# Powerful synergistic effects of a STING agonist and an IL-2 superkine in cancer immunotherapy against MHC I-deficient and MHC I+ tumors

**DOI:** 10.1101/2021.11.02.465022

**Authors:** Natalie Wolf, Cristina Blaj, Lora Picton, Gail Snyder, Li Zhang, Christopher J. Nicolai, Chudi O. Ndubaku, Sarah M. McWhirter, K. Christopher Garcia, David H. Raulet

## Abstract

Cyclic dinucleotides (CDNs) and TLR ligands mobilize antitumor responses by NK cells and T cells, potentially serving as complementary therapies to immune checkpoint therapy. In the clinic thus far, however, CDN therapy has yielded mixed results, perhaps because it initiates responses potently, but does not provide signals to sustain activation and proliferation of activated cytotoxic lymphocytes. To improve efficacy, we combined CDNs with a half-life extended IL-2 superkine, H9-MSA. CDN/H9-MSA therapy induced dramatic long-term remissions of the most difficult-to-treat MHC I-deficient and MHC I^+^ tumor transplant models. H9-MSA combined with CpG oligonucleotide also induced potent responses. Mechanistically, tumor elimination required CD8 T cells and not NK cells in the case of MHC I+ tumors and NK cells but not CD8 T cells in the case of MHC-deficient tumors. Furthermore, combination therapy resulted in more prolonged and more intense NK cell activation, cytotoxicity and expression of cytotoxic effector molecules in comparison to monotherapy. Remarkably, in a primary autochthonous sarcoma model that is refractory to PD-1 checkpoint therapy, the combination of CDN/H9-MSA combined with checkpoint therapy yielded long-term remissions in the majority of animals, mediated by T cells and NK cells. This novel combination therapy has potential to activate responses in tumors resistant to current therapies and prevent MHC I-loss accompanying acquired resistance of tumors to checkpoint therapy.

**One sentence summary:** Powerful immunotherapy effects mediated by the combination of innate agonists and superkine.

## Introduction

Great advances have been made in the immunotherapy field over the past decade, vastly improving patient outcomes ^1,2^. While many patients with malignancies show remissions, including long-term remissions, for most cancers the majority do not. Most of the approved therapies are predicated on amplifying CD8^+^ T cell responses, through blocking inhibitory “checkpoint” pathways ^1,3^. Many cancers, however, do not respond to checkpoint blockade and evade CD8^+^ T cell responses for numerous reasons. Some tumors lack neoantigens for T cell recognition ^4,5^. In addition, loss of MHC class I (MHC I) expression has emerged as a major mechanism of acquired resistance to checkpoint therapy ^6-10^. Therefore, mobilizing other antitumor effector cells with distinct recognition requirements, such as NK cells, is an attractive approach to counteract evasion of T cell responses.

Natural Killer (NK) cells are cytotoxic innate lymphocytes that can kill virus-infected and cancerous cells ^11-14^. NK cells recognize stressed-induced ligands on unhealthy cells ^15,16^ or the loss of MHC class I ^17-19^ through a combination of activating and inhibitory receptors, engagement of which determines whether a target cell meets the criteria for being killed ^20^. NK cells can also produce pro-inflammatory cytokines, such as interferon-*γ* (IFN*γ*) and tumor necrosis factor-*α* (TNF-*α*), which can augment immune responses in the tumor microenvironment and in some cases kill tumor cells directly ^21-23^. These features of NK cells make them desirable therapeutic targets for exerting direct tumor cell killing as well as enhancing other immune responses against tumors.

The cyclic guanosine monophosphate-adenosine monophosphate synthase–stimulator of interferon genes (cGAS-STING) pathway is an innate immune sensing pathway triggered by cytosolic DNA, that has shown promise in preclinical studies as a therapeutic target. Binding of cytosolic dsDNA stimulates the enzyme cGAS to catalyze the synthesis of the second messenger 2’,3’ cyclic guanosine monophosphate–adenosine monophosphate (cGAMP) ^24,25^. cGAMP binds and activates the ER membrane protein stimulator of interferon genes, STING, leading to activation of both IRF3 and NF-*κ*B pathways, and the abundant production of type I interferons and other inflammatory cytokines ^26^. Tumor cells often exhibit elevated levels of DNA damage, leading to cGAS activation and production of cGAMP, which can act directly on STING in the tumor cells or can be released from tumor cells to activate STING in neighboring normal cells, resulting in cytokine production and spontaneous antitumor immune responses ^27,28^. The magnitude of spontaneous antitumor responses are often quite weak, however, emphasizing the need for therapeutic intervention.

Synthetic cyclic di-nucleotides (CDNs) serve as potent STING agonists that, when injected intratumorally or systemically, superactivate antitumor T cell responses ^29,30^. Similarly, we recently demonstrated that intratumoral injections of CDNs induce robust NK cell-mediated rejection of six different established MHC I-deficient murine tumors, generated by CRISPR/Cas9 disruption of the *B2m* gene in each line ^31^. The CDN injections induced high levels of type I IFN, which was essential for NK cell activation, and acted directly on NK cells as well as indirectly by inducing trans-presentation of IL-15 by DCs. We observed permanent remissions of >50% of tumors in four of the six MHC I-deficient models studied. In two models, the rate of permanent remissions was much lower: ∼20% in the MC38-*B2m-/-* model and 0% in the B16-F10-*B2m-/-* model ^31^. At the same time, early phase clinical trials of STING agonists in patients have not yielded sustained clinical remissions ^32,33^. Clearly, there is a need for improvements in this immunotherapy approach.

Cytokines are powerful activators of immune cells. IL-2 family cytokines, including IL-15 and high dose IL-2, stimulate enhanced functional activity and greater survival of NK cells ^34^. Native IL-2 binds poorly to IL-2 receptors on most NK cells, which lack the IL-2R*α* chain.

However, a mutant form of IL-2, the superkine H9 (also known as super-2), was selected to bind with high affinity to the IL-2R*β/γ* complexes that NK cells express ^35^. H9 activates NK cells at much lower concentrations than IL-2, and was shown to stimulate antitumor responses in several mouse tumor models ^35^. Furthermore, whereas NK cells often become desensitized after infiltrating MHC I-low tumors, H9 administration reversed or delayed the onset of NK cell desensitization and prolonged the survival of mice with such tumors, in an NK cell-dependent fashion ^36^.

STING agonists provide a potent initial stimulus for NK cell responses but may not provide continual cytokine production to sustain the responses. Therefore, we tested the combination of a STING agonist with repeated administrations of the H9 superkine to determine whether the effective responses can be sustained. The results demonstrate remarkable efficacy of this combination therapy in both MHC I-deficient and MHC I^+^ tumor models.

## Results

### Half-life extended IL-2 superkine, H9, synergized with CDNs to induce rejection of MHC class I-deficient tumors

We previously tested CDN therapy in six different MHC I-deficient (*B2m-/-*) subcutaneous tumor lines by establishing the tumors to a size of approximately 50 mm^3^ followed by injection of the STING mixed-linkage (2′3′) RR cyclic diadenosine monophosphate (also known as ADU-S100, hereafter referred to as CDN) intratumorally. Poor efficacy was observed in two models, the B16-F10-*B2m-/-* melanoma model (no long-term survivors) and the MC38-*B2m-/-* colorectal cancer model (∼20% long-term survivors) showing the need for improvement. We therefore elected to test whether CDN therapy effects could be augmented with the H9 superkine. Cytokines such as IL-2 and H9 have very short half-lives *in vivo,* such that cytokine injections result in spikes of cytokine signaling followed by periods devoid of signaling ^37^. Fusing IL-2 to serum albumin has been shown to greatly extend its half-life, resulting in better efficacy and reduced toxicity ^37^. Therefore, we tested a fusion protein in which mouse serum albumin (MSA) was fused to the N-terminus of H9 (H9-MSA). Initially, we compared H9 and H9-MSA as monotherapies in the B16-F10-*B2m-/-* model in C57BL/6 (B6) mice. Once tumors were established s.c. with a high dose of tumor cells, they were treated every two or three days i.p. with H9 or H9-MSA. In cases where complete tumor remissions occurred, cytokine therapy was discontinued one week later. Compared to injections with PBS, injections of 5 *μ*g or 20 *μ*g H9-MSA slowed tumor growth and extended survival of the tumor bearing mice to a similar extent, whereas 20 *μ*g doses of H9 had no significant antitumor activity (Fig S1A).

In the same tumor model, a single i.t. injection of CDN also slowed tumor growth, but combining CDNs with 10 *μ*g or 20 *μ*g doses of H9-MSA provided dramatically improved outcomes, resulting in long-term (>100 d) tumor-free survival in 75-85% of the mice (Fig. 1A, B, Fig S1B, S2A). Efficacy fell off sharply with lower doses of H9-MSA (Fig. S1B). The combination of CDNs with H9, instead of H9-MSA, was significantly less efficacious, and CDNs combined with IL-2 showed almost no improvement in outcomes compared to CDNs alone (Fig. S1C).

**Figure 1.**
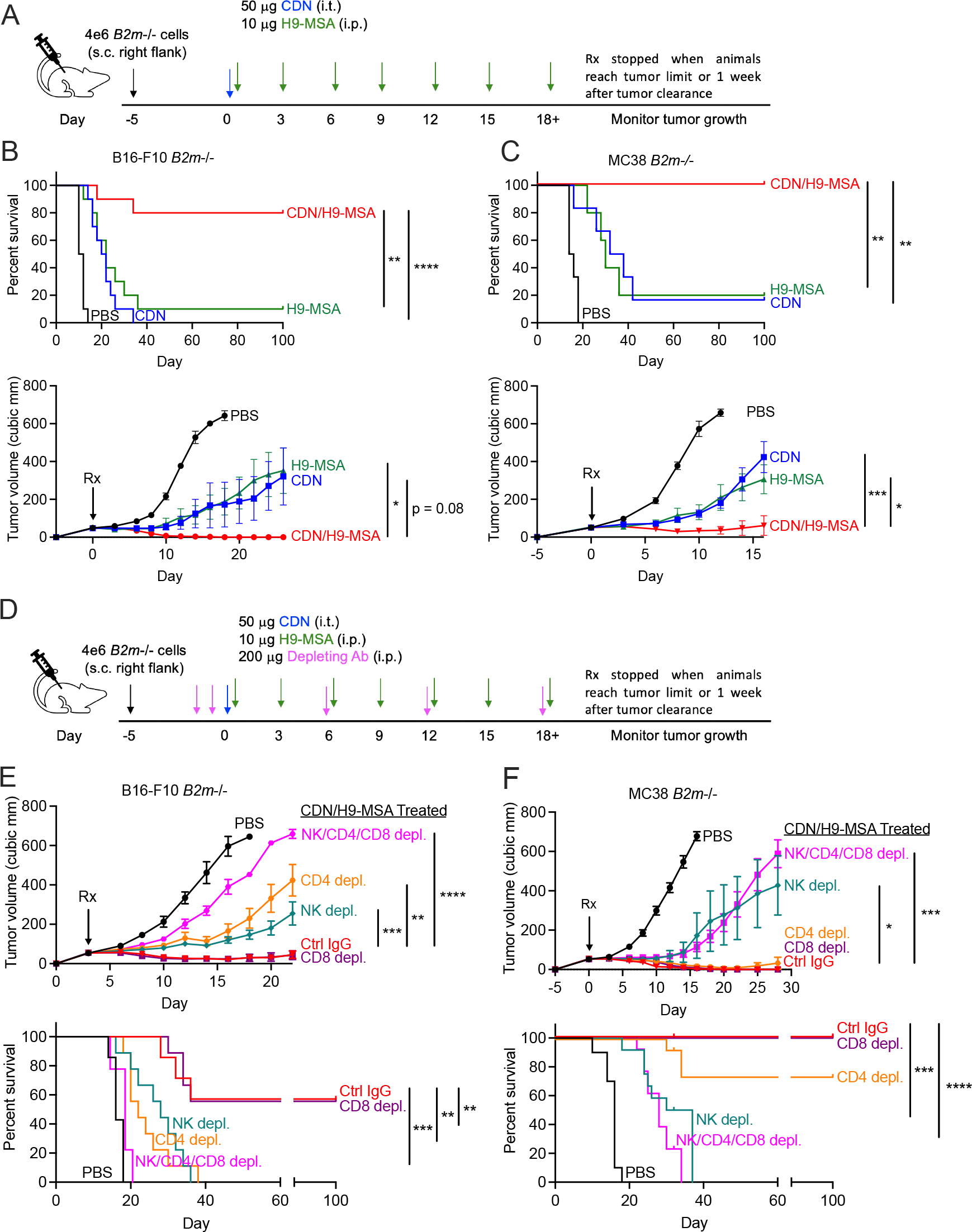
Rejection of MHC I-deficient tumors induced by CDN combined with IL-2 superkine, H9-MSA was mediated by NK cells and in one case by CD4 T cells. **(A)** Experimental timeline. Tumors were established with 4 x 10^6^ cells implanted s.c. in C57BL/6J mice and grown to approximately 50 mm^3^ (day 0). Tumors were injected once i.t. with 50 *μ*g of CDN or PBS. Mice were also injected i.p. with 10 *μ*g H9-MSA or PBS on day 0 and repeated every three days until euthanization or 1 week after complete tumor clearance. **(B,C)** Tumor growth curves and survival of mice with B16-F10-*B2m*-/- (B) and MC38-*B2m*-/- (C) tumors. **(D)** Experimental timeline for cellular depletions. Tumors were established and treated as in (A), except groups of mice were injected i.p. on days -2, -1 and every 6 days thereafter with antibodies to deplete NK cells, CD8 T cells and/or CD4 T cells, or with control IgG. **(E, F)** Tumor growth curves and survival of mice with B16-F10-*B2m*-/- (E) and MC38-*B2m*-/- (F) tumors. Tumor growth data was analyzed by 2-way ANOVA. Survival data was analyzed using log-rank (Mantel-Cox) tests. The data in all panels are representative at least two independent experiments. n=7-10 mice per group.

The CDN/H9-MSA combination gave even more impressive results in the MC38-*B2m-/-* model, resulting in long-term tumor-free remissions in 100% of the animals (Fig 1C, Fig. S2B).

In this model, CDN or H9-MSA treatments alone resulted in a small percentage of mice with long-term remissions and delays in tumor growth in the remaining animals (Fig. 1C, Fig. S2B). Thus, the combination of STING agonist and half-life extended H9 superkine was extremely effective in eliminating tumors in two very difficult-to-treat MHC I-deficient murine cancer models.

The combination of CDNs (one treatment) and 10 *μ*g H9-MSA (every two days) resulted in modest weight loss and a minor increase in the lung/body weight ratio, a metric for pulmonary edema, a known side effect of IL-2 therapy ^38^ (Fig. S3A-C). Greater weight loss and pulmonary edema were associated with the 20 *μ*g dose of H9-MSA by itself. CDNs alone had the least toxicity in all tests performed, corresponding to transient weight loss that in a separate experiment was maximal on day 1. Administering 10 *μ*g H9-MSA every three days further minimized toxicity while maintaining efficacy, and we employed that dose and schedule for most of the experiments in this study.

### The efficacy of CDN/H9-MSA combination therapy in MHC class I-deficient tumors was NK cell-dependent and CD8 T cell-independent

To identify effector cells responsible for rejection in mice receiving combination therapy, lymphocyte populations were depleted with repeated antibody treatments, starting after tumors were established but just before therapy was initiated (Fig 1D, Fig. S4). In the MC38-*B2m*-/- model, depleting either CD4+ or CD8*β*+ cells did not significantly diminish efficacy, whereas depleting NK cells alone greatly diminished therapeutic efficacy, resulting in terminal morbidity of all the animals (Fig 1F, Fig. S2D). Depleting all three types of lymphocytes had no greater effect than depleting NK cells, indicating that NK cells, and not CD4 or CD8 T cells, mediate tumor rejection.

In the B16-F10-*B2m*-/- model, depleting CD8 T cells had no effect, but depleting either NK cells or CD4 T cells diminished the efficacy of combination therapy (Fig 1E, Fig. S2C).

Depleting all three populations had an even greater effect, suggesting that combination therapy separately mobilized both antitumor NK cells and antitumor CD4 T cells, and that each acted to some extent independently. Antitumor CD4 T cells were previously implicated as effector cells in studies of wild type B16 tumors ^39,40^.

Some delay in tumor growth was still evident after depleting these three lymphocyte populations in both tumor models, which may reflect the impact of cytokines, including collapse of the tumor vasculature caused by TNF-*α* induced after CDN administration ^30,31,41^. Indeed, in the B16-*B2m-/-* tumor model, in B16-*B2m-/-* -bearing mice depleted of NK cells, CD4 T cells and CD8 T cells, simultaneous blockade of TNF-*α* and type I interferon receptors partially reversed the delay in tumor growth induced by CDN/H9-MSA administration (Fig. S5).

The NK1.1 antigen that is targeted by NK-depleting antibodies is also expressed by certain rare T cell populations. Furthermore, it remained possible that other nonconventional T cell populations that lack CD8 or CD4 participated in tumor rejection. To examine whether these other cell types might play a role, we tested the combination therapy in tumor-bearing B6-*Rag2-/-* mice, which lack T cells (including NK1.1^+^ T cells) and B cells but retain NK cells. In both tumor models, H9-MSA and CDN treatments were more potent in combination than separately in the B6-*Rag2-/-* mice (Fig 2). In the MC38-*B2m*-/- model, the combination therapy was as effective in *Rag2-/-* mice as in WT mice, resulting in 100% survival of the mice (Fig. 2A, C, Fig. S6A). As before, NK-depletion greatly diminished efficacy, resulting in 100% mortality. In the B16-F10-*B2m*-/- model, consistent with the effects of CD4 depletion in WT mice, combination therapy only delayed tumor growth in *Rag2-/-* mice, and 100% of the animals eventually succumbed (Fig 2B, D, Fig. S6B). NK depletion in *Rag2-/-* mice accelerated tumor growth, as expected. Thus, combination therapy mobilized a fully effective antitumor NK response against MC38-*B2m-/-* tumors, with no discernable role for T cells, whereas both antitumor NK cells and antitumor CD4 T cells acted in parallel, albeit to a significant extent independently, to eliminate B16-*B2m-/-* tumors in WT mice. CD8 T cells played no discernable role in either response.

**Figure 2.**
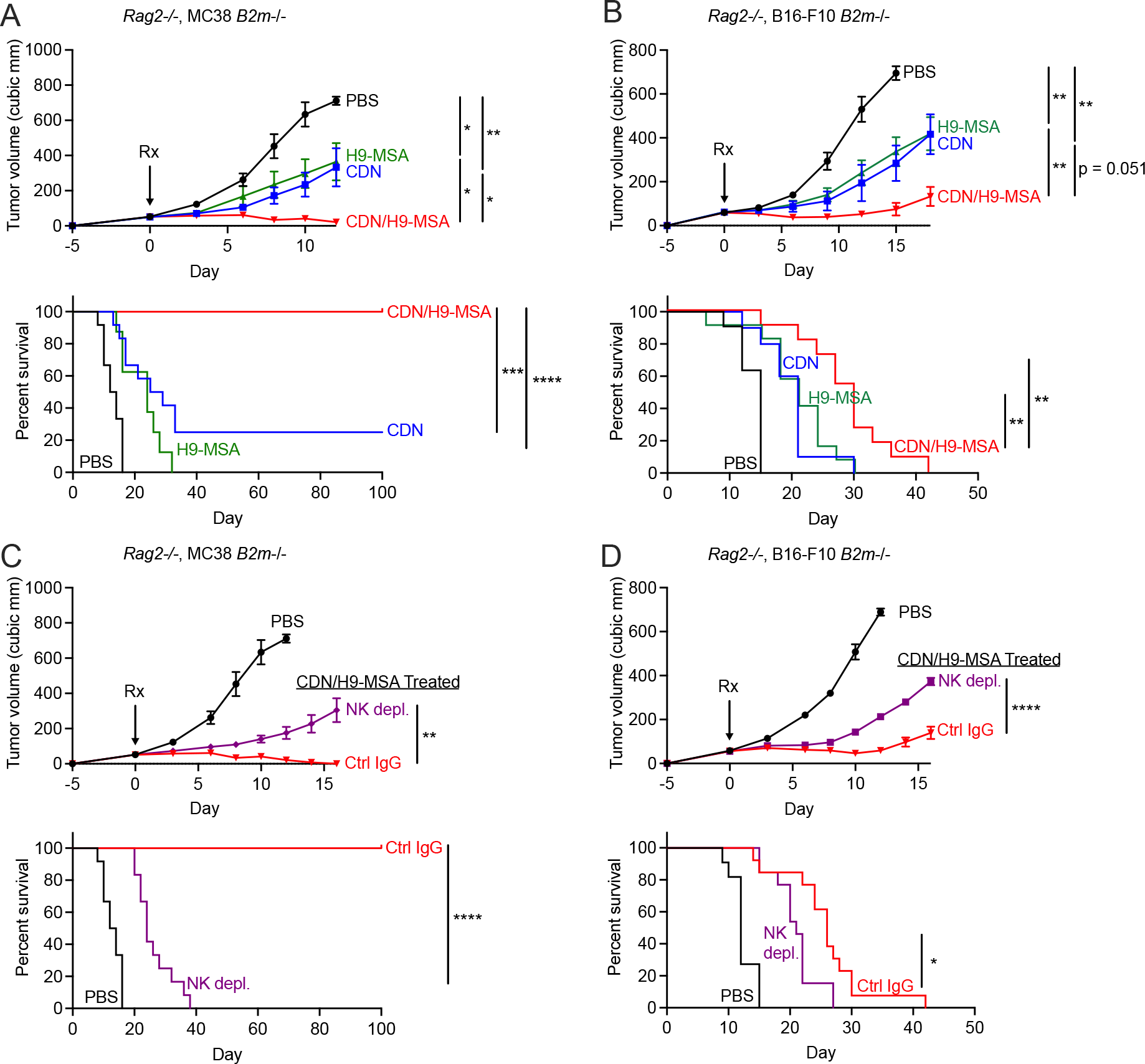
NK cell dependent rejection of MHC I-deficient tumors induced by CDN/H9- MSA treatment in *Rag2-/-* mice. MC38-*B2m*-/- (A, C) and B16-F10-*B2m*-/- (B, D) tumors were established as in Fig. 1 legend in *Rag2*-/- mice, and subjected to therapy as indicated with CDN, H9-MSA, the combination as in Fig. 1 legend. **(A, B)** Combination therapy is more effective than monotherapies in both tumor models. **(C, D)** The therapeutic effects were decreased by NK-depletion in both models. Tumor growth data were analyzed by 2-way ANOVA. Survival data were analyzed using log-rank (Mantel-Cox) tests. The data in all panels are representative at least two independent experiments. n=9-12 mice per group.

### H9-MSA was more effective than WT IL-2-MSA in combination with CDNs

We next compared H9-MSA directly with WT IL-2-MSA in the context of the combination therapy in WT mice with established B16-*B2m-/-* tumors. The mice were depleted of CD4 and CD8 T cells before the initiation of therapy to narrow the analysis to the effects of NK cells. The CDN/H9-MSA combination was again more effective CDN or H9-MSA alone (Fig. 3A). The CDN/IL-2-MSA combination, in contrast, was only slightly more effective than CDN or IL-2-MSA alone (Fig. 3B). In the head-to-head comparison, the CDN/H9-MSA combination was more effective than the CDN/IL-2-MSA combination, both in terms of delaying tumor growth and delaying terminal morbidity (Fig. 3C, Fig. S7A). Therefore, H9- MSA is more effective than IL-2-MSA in activating NK cells against B16-*B2m-/-* tumors.

**Figure 3.**
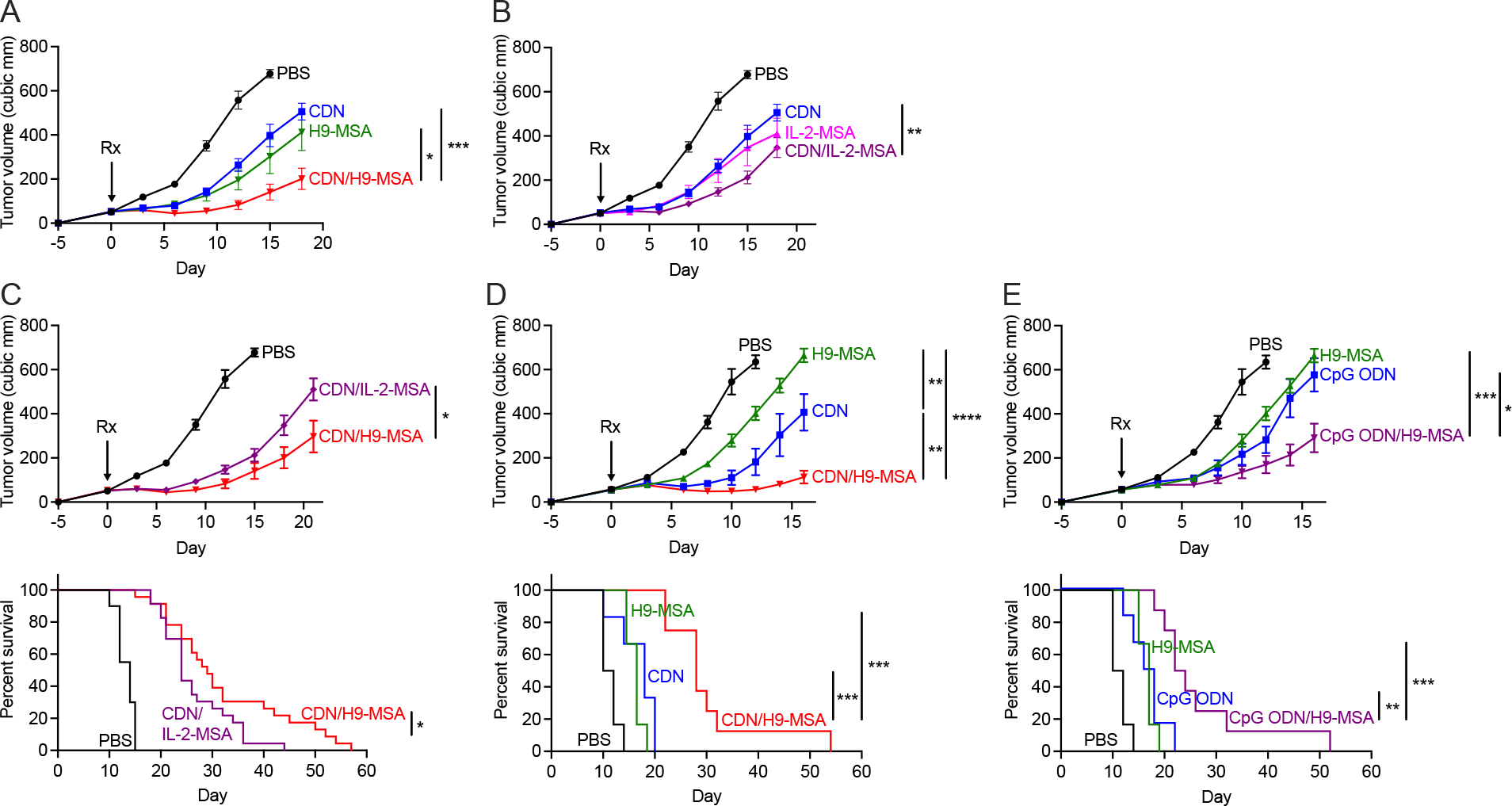
Greater efficacy of H9-MSA compared to IL-2-MSA in mice lacking CD4 and CD8 T cells, and synergy of CpG ODN and H9 MSA. B16-F10-*B2m*-/- tumors were established as in Fig. 1 legend in C57BL/6J mice. In all panels, the mice were depleted of both CD4 and CD8 T cells before initiating therapy. **(A-C)** Comparisons of the effects of CDN, H9-MSA and IL-2-MSA separately (A, B) or in combinations (CDN/IL- 2-MSA vs CDN/H9-MSA) (A-C) using the schedule shown in Fig. 1, one dose of 50 *μ*g of CDN and 10 *μ*g doses of the cytokine fusion proteins. Curves in A-C were all from the same experiment. **(D, E)** Comparison of the effects of CDN, CpG ODN (each at 50 *μ*g dose) and H9- MSA (10 *μ*g doses) separately or in combinations (CDN/H9-MSA vs CpG ODN/H9-MSA). Curves in D and E were from the same experiment. Tumor growth data were analyzed by 2-way ANOVA. Survival data were analyzed using log-rank (Mantel-Cox) test. The data in all panels are representative at least two independent experiments. n=9-20 mice per group.

### A distinct innate activator, CpG oligonucleotide, also synergized with H9-MSA in providing antitumor efficacy

Aside from CDNs, other innate agonists are known to stimulate production of interferons and other cytokines downstream of IRF3 and NF-*κ*B. One such agonist is the TLR9 ligand CpG oligodeoxynucleotides (ODN). We therefore compared i.t. injections of CpG ODN and CDNs in the context of combination therapy in WT mice with established B16-*B2m-/-* tumors. Again, the mice were depleted of CD4 and CD8 T cells before the initiation of therapy. Similar to CDNs, CpG ODN by itself delayed tumor growth and synergized with H9-MSA to induce more potent antitumor effects, though not quite to the same extent as did CDNs at the same dose level (Fig. 3D, E, Fig S7B). These findings indicated that there is some generality to the synergistic effects resulting from combining H9-MSA with TLR agonists, CDNs or possibly other stimulants of the innate immune system.

### Local CDNs and systemic H9-MSA synergized in creating a systemic NK cell mediated antitumor effect

When tumors metastasize, treatment options are more limited and lead to greater patient mortality. To address the impact of CDN/H9-MSA therapy on creating a systemic NK cell effect with the potential to reject metastases, we established a two-tumor model system. B16-F10-*B2m*-/- tumors were established on both flanks of syngeneic mice (Fig. 4A). In order to focus on the NK-mediated effect, all the mice were depleted of CD4 and CD8 T cells before the start of treatment. Tumors on the right flank of each mouse were injected with either PBS or CDN. H9- MSA or PBS was then delivered intraperitoneally. As before, combination therapy was more effective than the single therapies in the CDN-injected tumors (Fig. 4B, Fig. S8A). Remarkably, the same was true in the tumors that were not injected with CDNs, corroborating that local CDN therapy by itself exerted a systemic effect ^31^, and demonstrating that CDNs synergized with H9- MSA in mobilizing greater systemic antitumor effects (Fig. 4B, Fig. S8A). Depletion of NK cells resulted in diminished efficacy of the combination therapy similarly in both the injected tumor and the uninjected tumor (Fig. 4C, Fig. S8). As these mice were depleted of CD4 and CD8 T cells, the data implicated NK cells as major antitumor effectors both locally and in the distant tumor.

**Figure 4.**
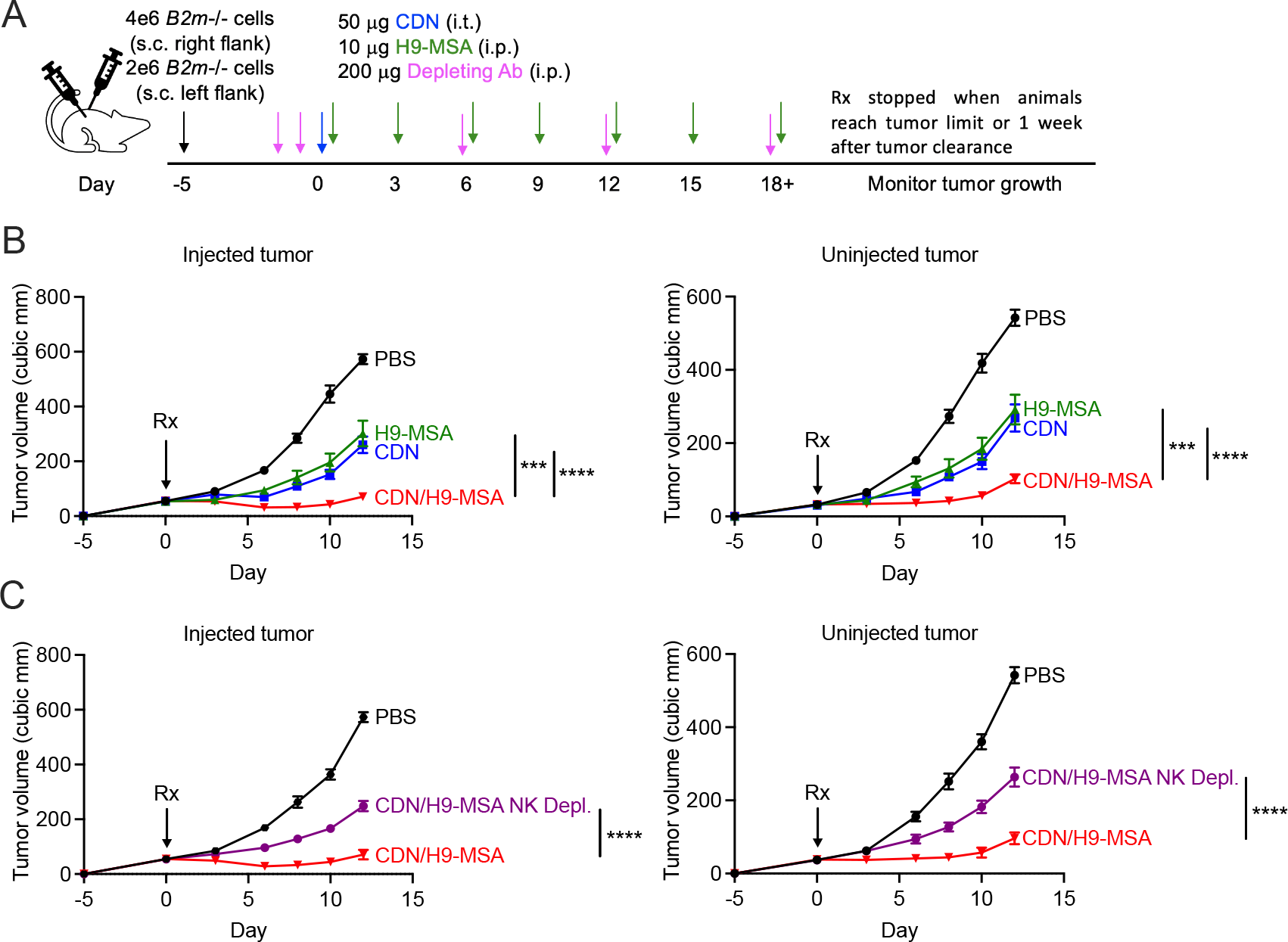
Intratumoral CDN injections combined with intraperitoneal H9-MSA injections synergized in creating a systemic NK cell-mediated antitumor effect. **(A)** Timeline of the two-tumor experiments. On day -5, B16-F10-*B2m*-/- tumor cells were injected s.c. on both flanks of C57BL/6J mice, with 4 x 10^6^ cells on the right flank and 2 x 10^6^ cell on the left flank. Mice were depleted of CD4 cells and CD8 cells 2 days and 1 day before initiation of treatment. On day 0, the tumor on the right flank was injected once i.t. with 50 *μ*g CDN or PBS. The mice received i.p. injections of PBS or 10 *μ*g H9-MSA starting on day 0 and again every three days until mice were euthanized. **(B)** Growth of injected (left panel) vs contralateral uninjected tumors (right panel) receiving the indicated therapies. The data were combined from 3 experiments (n=18). **(C)** Tumor growth following CDN/H9-MSA therapy in mice from which NK cells were depleted or not (with the treatment schedule shown in panel A) as indicated. The data were combined from 2 experiments (n=12). Tumor growth data were analyzed by 2-way ANOVA.

### CDN/H9-MSA combination therapy mobilized more activated intratumoral NK cells with greater functional activity

To address the impact of CDN/H9-MSA treatment on NK cells, we employed the two-tumor system, and examined markers of activation on NK cells infiltrating both tumors 2 days and 5 days after treatment. On both days, NK cells in both the injected and uninjected tumors contained the most elevated percentages of granzyme B^+^, and the most elevated levels of granzyme B in the positive cells (Fig. 5A, B, Fig. S10, Fig. S11). In numerous cases, the differences between single treatments and combined treatments were statistically significant. Similarly, the percentages of NK cells expressing the activation markers CD69 or Sca1 were elevated in the injected and the uninjected tumors on both days (Fig. 5A, B, Fig. S9A, B). Again, the differences were in some cases statistically significant. NK cell percentages among CD45+ cells, and the fraction of the NK cells expressing the proliferative marker Ki67, also trended higher in some groups of mice receiving one or the other therapy, or both therapies (Fig. 5A, B, Fig. S10, Fig. S11).

**Figure 5.**
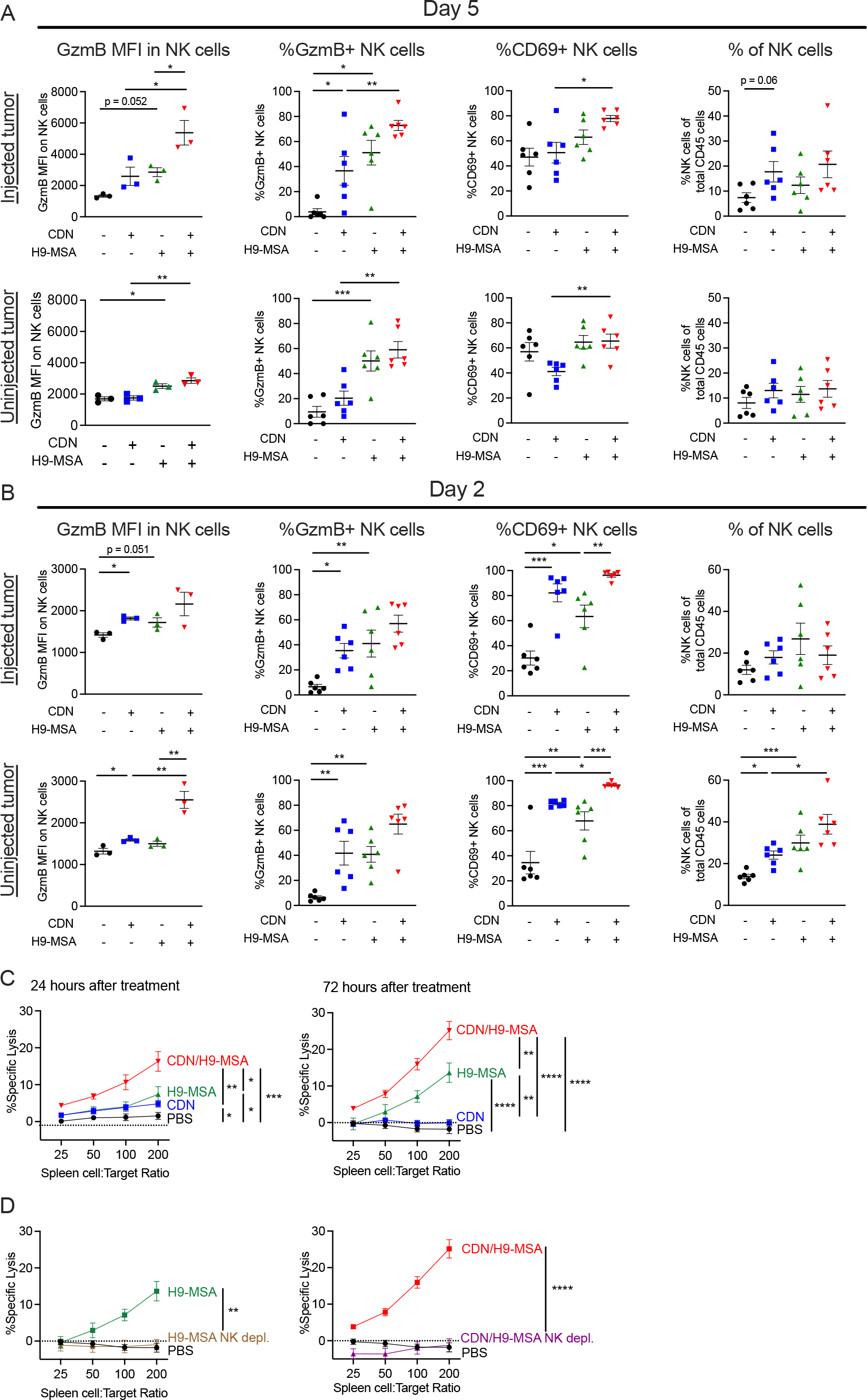
Activation, proliferation and cytotoxicity of NK cells in tumor bearing mice receiving CDN/H9-MSA immunotherapy. B16-F10-*B2m*-/- tumors were established on both flanks of C57BL/6J mice as in Figure 4. On d5 **(A)** or d2 **(B)** Single cell suspensions of both tumors were examined for the intracellular or cell surface markers shown and the percentages of NK cells among viable CD45+ cells. n=3. GzmB MFI data are representative of at least 2 independent experiments (n=3) and all other data were combined from 2 independent experiments (n=6). Samples were analyzed by one-way ANOVA. **(C)** B16-F10-*B2m*-/- tumors were established and treated as described in Figure 4. 24 hr and 72 hr after treatment, splenocytes were tested for cytotoxicity of B16-F10-*B2m*-/- target cells. Spontaneous lysis of these target cells averaged 11%. Error bars are shown for biological replicates (n=8). **(D)** B16-F10-*B2m*-/- tumors were established, treated and tested as in panel C, 72 hrs after initiating H9-MSA therapy or combination therapy, as indicated. Additionally, mice in one group receiving each therapy were depleted of NK cells by injecting 200 *μ*g NK1.1 antibody i.p. 2 days and 1 day before initiating therapy. n=8 biological replicates. Data in panels C and D were combined from 2 independent experiments and were analyzed by 2-way ANOVA. n=8.

To assess whether combination therapy induced NK cytotoxic activity systemically, splenocytes from tumor-bearing mice were tested for killing of B16-F10-*B2m*-/- tumor cells ex vivo 24 or 72 hours after initiating treatment (Fig. 5C). Some elevation of cytotoxicity was observed with H9-MSA or CDN treatments alone, but the killing with CDNs alone diminished by 72 hours. Significantly greater cytotoxicity was observed with splenocytes from mice receiving combination therapy. When NK cells were depleted from mice prior to initiating treatment, cytotoxicity was eliminated in the subsequent ex vivo assay, confirming that NK cells mediated the killing (Fig. 5D). These data demonstrated that the combination therapy had multiple impacts, maximizing NK cell activation and content of cytotoxic effector molecules (granzyme B), as well as cytotoxicity, and prolonging the NK response compared to each therapy separately.

### CDN/H9-MSA therapy mobilized CD8 T cells against MHC I^+^ tumors

The impact of combination therapy on NK cell-dependent tumor rejection prompted us to ask whether this approach has broader applicability, including against MHC I^+^ tumors.

Therefore, we tested whether the CDN/H9-MSA therapy combination is effective with “wildtype” MHC I^+^ B16-F10 tumors, a model that is poorly responsive to most therapies including checkpoint therapy. B6 mice with established MHC I^+^ B16-F10 tumors were treated with CDNs and/or H9-MSA using the same scheme as for MHC I-deficient tumor (Figure 1A). Mice treated separately with CDNs or H9-MSA showed delays in tumor growth, but very poor long-term survival, whereas the combination of CDN/H9-MSA showed synergistic tumor rejection and disease-free survival of most of the mice for > 100 days (Fig. 6A, Fig. S12A).

**Figure 6.**
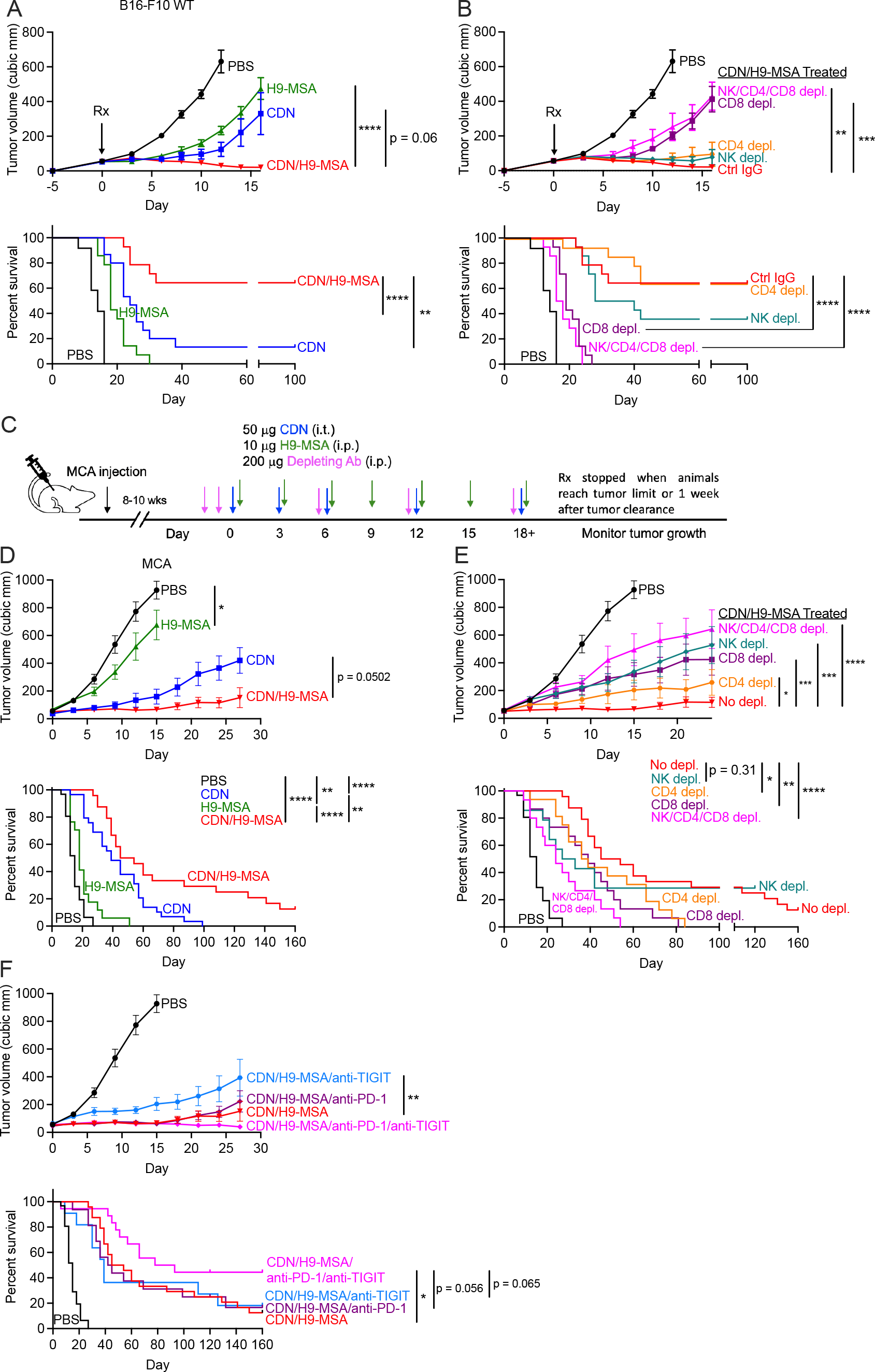
CDN/H9-MSA therapy induced potent antitumor effects with MHC I+ tumors, mediated by CD8 T cells. **(A, B)** Experiments were carried out as in Fig. 1 except MHC I+ B16-F10 wildtype cells were employed. **(A)** Impact of CDN, H9-MSA or combination therapy. **(B)** Antitumor effects were reversed by depleting CD8 T cells (p<0.0001), with no significant impacts of depleting NK cells or CD4 T cells (p=ns). Tumor growth data were analyzed by 2-way ANOVA. Survival data were analyzed using log-rank (Mantel-Cox) test. The data in all panels are representative of two independent experiments. n=6-7 mice per group. **(C)** Experimental timeline for MCA-induced tumors. When MCA-induced tumors reached ∼50 mm^3^ in size they were injected i.t. with PBS or 50 *μ*g CDN on days 0, 3, and 6 and repeated every 6 six days, thereafter. Mice were also injected i.p. with PBS or 10 *μ*g H9-MSA starting on day 0, which was repeated every three days. In some groups, mice were injected i.p. on days -2, -1 and every 6 days thereafter with antibodies to deplete NK cells, CD8 T cells, CD4 T cells, or with control IgG. **(D)** Tumor growth curves and survival of mice. **(E)** Tumor growth curves and survival of depleted mice. **(F)** Addition of checkpoint inhibitors consisting of i.p. injections of anti-PD-1 (clone RMP1-14) and anti-TIGIT (clone MUR10A), increased tumor rejection and overall survival. Data in these panels are a combination of multiple experiments, such that some of the data (e.g. CDN/H9-MSA) are shared in different panels. n=11-24. Data were analyzed by 2-way ANOVA.

Cell depletion experiments demonstrated that CD8 T cells were essential for rejection of the MHC I^+^ tumors in the context of CDN/H9-MSA therapy, whereas CD4 cells and NK cells were largely dispensable (Fig 6B, Fig. S12B). Elimination of all three populations was no more impactful than CD8 depletion alone. Thus, CD8 T cells are very effectively mobilized against MHC I^+^ tumors by this combination therapy. Together with the MHC I-deficient tumor findings, the results demonstrated that combining innate agonists and IL-2 superkines synergistically mobilized T cells or NK cells against tumors, depending on the MHC expression pattern of the tumor cells.

### Efficacy of combination therapy in the methylcholanthrene induced sarcoma model

To address the efficacy of the CDN/H9-MSA combination in a primary autochthonous cancer model, we chose the methylcholanthrene (MCA)-induced sarcoma model. MCA was injected s.c. on the right flanks and mice were monitored frequently for palpable tumors, which arose 8-10 weeks after carcinogen administration in 80% of the treated animals. When a tumor reached a volume of approximately 50 mm^3^, the mouse was treated individually. Mice were initially injected with three doses of CDNs i.t. every three days followed by weekly injections. Some mice received H9-MSA i.p. applied every three days (Fig. 6C). H9-MSA alone had a minor, albeit significant effect on tumor growth and terminal morbidity of the animals, whereas CDN therapy caused a substantial delay in both tumor progression and terminal morbidity (Fig. 6D, Fig. S13A). The CDN/H9-MSA combination was significantly more effective than CDNs alone in prolonging survival, with a small percentage of animals surviving after 160 days.

Some toxicity of the CDN/H9-MSA combination was evident in this model, including the appearance of chronic granulomatous histiocytic inflammation in some mice, that arose in the leg muscle on the same side of the mouse as the treated tumor. When treated with topical diphenhydramine (Benadryl), the inflammation was partially controlled, and mice were continued in the experiment.

The initial antitumor effects were significantly diminished when either NK cells, CD8 cells or, to a lesser extent, CD4 cells were depleted from the animals before initiating therapy (Fig. 6E, Fig. S13B). Depleting NK cells accelerated the demise of a fraction of the animals, whereas depleting CD8 or CD4 cells accelerated the death of all the animals. Depleting all three populations simultaneously had a modestly greater effect than depleting CD8 or CD4 cells separately, suggesting some separate impacts of these lymphocyte populations, though some cooperative interactions between the populations are fully consistent with the data. Thus, NK cells, CD8 T cells and CD4 T cells all contribute to the therapeutic effects in this tumor model.

Although the CDN/H9-MSA combination showed efficacy in treating primary MCA tumors, nearly all the mice eventually succumbed. Therefore, we tested whether efficacy could be improved by adding checkpoint therapy to the combination. Addition of anti-PD-1 therapy did not result in an improvement in survival, nor did addition of anti-TIGIT therapy (Fig. 6F, Fig. S13C). However, simultaneous blockade of PD-1 and TIGIT, in combination with CDN/H9- MSA, was significantly more effective than CDN/H9-MSA therapy alone (Fig. 6F, Fig. S13C). Indeed, 44% of the animals (8/18) receiving the four-component therapy regimen survived tumor-free for >160 days. Interestingly, the chronic inflammation noted in some mice treated with CDN/H9-MSA was not observed in the mice receiving all four agents. Some cases of severe acute toxicity were observed when the four components were injected simultaneously, but delaying the anti-TIGIT treatments by one day appeared to ameliorate that toxicity. Mice that received anti-PD-1 or anti-TIGIT treatments alone or together did not show prolonged survival (Fig. S13D). CDN therapy outcomes were also not improved by the addition of anti-PD-1 or anti-TIGIT separately (Fig. S13E). In conclusion, simultaneous blockade of PD-1 and TIGIT was necessary to unleash a fully effective antitumor immune response initiated by CDN/H9- MSA treatments.

## Discussion

The CDN/H9-MSA therapy combination induced profound antitumor responses in several solid tumor models. A single intratumoral injection of CDNs combined with periodic injections of the IL-2 superkine, H9-MSA, resulted in a dramatically increased percentage of animals with disease-free survival in three very difficult-to-treat tumor transplant models, two of which were MHC I-deficient and refractory to CD8 T cell recognition. In primary MCA-induced sarcomas, the combination therapy, by itself, delayed tumor growth, but led to disease-free survival in many animals when combined with immune checkpoint therapy. Depending on the tumor model, tumor rejection depended on NK cells, both NK cells and CD4 T cells, CD8 T cells, or all three populations. The long-term remissions that we have documented and the efficacy in mobilizing both T cells and NK cells, suggest that combinations of this type should be considered for testing in human cancer patients. Initial tests may be especially warranted in instances where acquired resistance to checkpoint therapy has occurred.

High dose IL-2 has been approved as a monotherapy for metastatic renal carcinoma and melanoma leading to tumor regressions in a small subset of patients ^42^. It is only effective in a few patients, however, and is associated with vascular leak syndrome, heart failure and liver toxicity, and can favor the activation of regulatory T cells with the potential to counteract its immunostimulatory effects. H9 was previously shown to mobilize fewer regulatory T cells and to be less toxic than IL-2 ^35^, and we observed acceptable toxicity with effective doses of H9-MSA in mice. Published studies show greater efficacy of IL-2 and other cytokines with half-life extending modifications, such as fusion to albumin or Fc domains, or PEGylation ^37,43^. We observed that extended half-life H9 was more effective than extended half-life IL-2 in the context of combination therapy. Considering these results, H9-MSA warrants consideration for clinical applications. Nevertheless, efficacy may also occur when combining CDN therapy with other half-life extended IL-2 and IL-15 variants, several of which are undergoing clinical testing^43^.

In syngeneic tumor transplant models, H9-MSA acted synergistically not only with CDNs, but also with another agonist for the innate immune system, CpG ODN DS-L03, a TLR9 agonist optimized to stimulate high levels of type I IFN ^44^. These findings show the promise of combining IL-2/IL-15 family cytokines with agents that induce IRF3 and NF-*κ*B signaling.

Clinical trials of TLR agonists and STING agonists as cancer therapies have thus far yielded mixed results but clinical development remains active ^32,33,45^, and our results suggest that combining those agents with cytokines, such as H9-MSA, may have considerable promise.

The synergistic antitumor effects of CDNs and H9-MSA are likely due in part to the potent effects of IL-2 family cytokines on NK cells, including stimulating proliferation and activation of effector functions ^46^ and reversing or preventing NK cell desensitization that occurs in MHC I-deficient tumors ^47^. Intratumoral injections of CDNs induce significant levels of type I IFN systemically, which potently activates NK cells directly and indirectly ^31^. The direct effects include induction of effector molecules, such as granzymes ^31,48^, and protection from NK cell fratricide ^49^. The indirect effects are mediated in part by induction of IL-15R*α*/IL-15 complexes primarily on DCs, which are trans-presented to NK cells ^31,50^. The transient nature of CDN therapy, however, means that the early burst of IFN and IL-15 expression are not sustained. In the absence of sustained availability of IL-15 and other cytokines, primed NK cells infiltrating MHC class I-deficient tumors may more rapidly acquire an anergic state, associated with ineffective signaling and degranulation and overall reduced tumor cell killing. In mice with MHC I-deficient tumors, cytokines, such as H9, or a mixture of IL-12 and IL-18, have been shown to reverse or inhibit NK cell anergy, slowing tumor growth and extending survival ^47^.

Consistent with these considerations, mice treated with the CDN/H9-MSA combination showed elevated expression of activation markers and granzyme B among tumor-infiltrating NK cells, even in tumors distant from the site of CDN administration, and sustained cytolytic activity of splenic NK cells against MHC I-deficient tumor cells.

The CDN/H9-MSA combination therapy mobilized different effector cells depending on the tumor cell line studied and whether the tumor cells expressed MHC I molecules. First, the combination therapy induced NK cell mediated antitumor responses in two different MHC I- deficient tumor models. Remarkably, in the difficult-to-treat MC38-*B2m*-/- model, the treatment effectively cured 100% of tumor-bearing *Rag2*-/- mice, which lack T cells. NK cells were essential for these dramatic effects.

In the B16-F10-*B2m*-/- model, the combination therapy mobilized both antitumor NK cells and antitumor CD4 T cells, which were sufficient together, but not separately, to eliminate tumors and lead to long-term survival of the large majority of animals. The less effective NK cell responses in the B16-F10-*B2m*-/- model could be related to the absence of NKG2D ligands, which are present on MC38 cells ^16^. The elicitation of antitumor CD4 T cells in the B16-F10- *B2m-/-* model, but not the MC38-*B2m-/-* model, might be explained by the finding that MHC II expression can be induced on B16-F10 cells in vivo, but other features of the tumor cells may also be at play given that antitumor CD4 T cells have been described for B16 tumor cells in several other scenarios ^39,51,52^.

In the context of MHC I+ B16-F10 tumors, considered a “cold” tumor model that is resistant to checkpoint therapy and other experimental monotherapies ^53^, CDN/H9-MSA therapy induced a highly effective CD8 T cell mediated antitumor response, with no evidence for roles of NK cells or CD4 T cells. The impressive impact of CDN/H9-MSA therapy shown here is notable considering that durable antitumor responses in this model has required combining 3-4 agents in past studies ^30,54^. The absence of effective antitumor NK cells in this model with CDN/H9-MSA therapy can be explained by the inhibitory effects of MHC I, but it remains unclear why effective antitumor CD4 T cells were not elicited.

The finding that CDN/H9-MSA therapy worked well for both MHC I+ cold tumors and MHC I-deficient tumor variants suggests the promise of this approach both for tumor types that are resistant to checkpoint therapy and for preventing the emergence of MHC I-deficient tumor variants during immunotherapy. The loss of MHC I expression may occur before or during immunotherapy and is increasingly recognized as a significant mechanism of acquired resistance of tumors to checkpoint therapy ^8-10,55,56^.

CDNs and H9-MSA also synergized in the MCA sarcoma model, leading to highly delayed tumor growth and extended survival. Both NK cells and T cells contributed to the therapeutic effect, providing a demonstration that this therapy combination can mobilize both types of responses in the same tumor model. The addition of checkpoint therapy to the CDN/H9- MSA therapy regimen resulted in tumor regressions and long-term tumor free survival in approximately 44% of the treated animals. These findings represent the first example of long-term immunotherapy remissions in an autochthonous sarcoma model in mice, which is an especially substantive outcome considering that immunotherapy has given generally poor outcomes in treating human sarcomas ^57^.

In the absence of T cells and NK cells, the combination therapy still delayed tumor growth in all the tumor models studied but did not result in long-term remissions. Our data suggest that therapy-induced production of TNF-*α*, which alters the tumor vasculature, and type I IFNs, which inhibit cell proliferation, are partly responsible for the residual antitumor effects. It remains possible that the therapy combination also induces other cell types that retard tumor growth, such as myeloid cells or other innate lymphoid cell subtypes.

An important consideration for clinical applications is whether therapies elicit control of metastases distant from the site of a primary tumor, which are known to limit the efficacy of immunotherapies ^58,59^ We observed that local administration of CDNs in a primary tumor, combined with systemic applications of H9-MSA, resulted in strong systemic activation of NK cells capable of limiting the growth of a second tumor on the opposite flank of the animals and mediating killing of tumor cells in ex vivo tests with splenic NK cells. In contrast, NK cell activation after a single treatment with CDNs alone was short lived, as by 72 hours after treatment, cytotoxicity by splenic NK cells had subsided. Thus, the combination therapy sustained the systemic cytotoxicity of NK cells. Furthermore, in both the injected and uninjected tumors, the combination therapy mobilized significant increases in NK cell effector molecule expression and activation markers that were sustained over time compared to either CDNs or H9- MSA alone. The impact of intratumoral injections of CDNs on NK cell activation in a distant second tumor or in the spleen may be due to the burst of systemic cytokines induced by intratumoral CDN injections ^31^ and/or the leakage of small amounts of CDNs from the tumor into the general circulation ^30^.

In conclusion, our results show that CDN therapy combined with the IL-2 superkine, H9- MSA, effectively enhanced the rejection of MHC I+ and MHC I-deficient tumors, as well as primary carcinogen induced sarcomas. The CDN/H9-MSA treatments proved to be remarkably effective against difficult-to-treat tumors, mobilizing various effector cells depending on tumor cell type and MHC I expression. In MHC I-deficient tumors, the combination therapy elicited substantial systemic NK cell activity and antitumor effects on tumors distant from the injected tumors. These results provide compelling support for testing combinations of innate immune system agonists and IL-2 family superkines as potential next-generation immunotherapies for tumors that are resistant to currently approved immunotherapy regimens.

## Materials and Methods

### Study design

For most experiments in this study, tumors were established subcutaneously in mice, PBS or CDNs were injected intratumorally once, and PBS or H9-MSA were injected intraperitoneally multiple times. Tumor growth, toxicity, overall survival, and NK cell activation status were recorded. Male and female mice were used equally, and treatments were randomized among mice in the same cage. Experimental groups consisted of 5-10 mice. Mice were terminated in the studies when tumors reached an average diameter of 1.5 cm or mice showed a body condition score less than 2. We did not use a power analysis to calculate sample size. All experiments were performed at least twice, and in some cases, experiments were pooled. The investigators were not blinded.

### Mouse strains

Mice were maintained at the University of California, Berkeley. C57BL/6J and *Rag2-/-* mice were purchased from the Jackson Laboratory. All mice used were between 8 and 20 weeks of age. All experiments were approved by the University of California (UC) Berkeley Animal Care and Use Committee and were performed in adherence to the NIH Guide for the Care and Use of Laboratory Animals.

### Cell lines and culture conditions

B16-F10 cells (obtained from the UC Berkeley Cell Culture Facility) and MC38 cells (obtained from JP Allison) were cultured in Dulbecco’s modified Eagle’s medium (ThermoFisher Scientific). *B2m-/-* versions of both B16-F10 and MC38 were previously generated in the lab using CRISPR-Cas9 ^31^. In all cases, medium contained 5% fetal bovine serum (FBS) (Omega Scientific), 0.2 mg/mL glutamine (Sigma-Aldrich), 100 U/mL penicillin (ThermoFisher Scientific), 100 *μ*g/mL streptomycin (ThermoFisher Scientific), 10 *μ*g/mL gentamycin sulfate (ThermoFisher Scientific), 50 *μ*M *β*-mercaptoethanol (EMD Biosciences), and 20 mM Hepes (ThermoFisher Scientific). Cells were cultured in 5% CO2. All cell lines tested negative for mycoplasma contamination.

### Protein Design and Purification

DNA encoding wild-type human IL-2 or human IL-2 with H9 mutations (L80F, R81D, L85V, I86V, I92F) (*35*) was cloned into the insect expression vector pAcGP67-A, which includes a C-terminal 8xHIS tag for affinity purification. DNA encoding mouse serum albumin (MSA) was purchased from Integrated DNA Technologies (IDT) and cloned into pAcGP67-A as an N-terminal fusion between the N-terminus of hIL-2 and C-terminus of MSA (indicated as IL-2- MSA or H9-MSA).

Insect expression DNA constructs were transfected into Trichoplusia ni (High Five) cells (Invitrogen) using the BaculoGold baculovirus expression system (BD Biosciences) for secretion and purified from the clarified supernatant via Ni-NTA followed by size exclusion chromatography with a Superdex-200 column and formulated in sterile Phosphate Buffer Saline (PBS) (Gibco). Endotoxin was removed using the Proteus NoEndo HC Spin column kit following the manufacturer’s recommendations (VivaProducts) and removal was confirmed using the Pierce LAL Chromogenic Endoxotin Quantification Kit (ThermoFisher). IL-2 proteins were concentrated and stored at -80 °C until use.

### In vivo tumor growth experiments

Cells were washed and resuspended in PBS (ThermoFisher Scientific). 100 *μ*l containing 4 × 10^6^ cells were injected s.c.. Tumor volume was estimated using the ellipsoid formula: V = (4/3)*π*abc where a, b and c correspond to height, width and length of the tumors measured with digital calipers. Five days after tumor cell inoculation, when tumors reached a volume of 50 mm^3^, they were injected i.t. with PBS or 50 *μ*g of the STING agonist mixed-linkage (2′3′) RR cyclic diadenosine monophosphate (ADU-S100, abbreviated as CDN in this paper, a gift of Aduro Biotech) in a total volume of 100 *μ*l PBS. Mice were also injected i.p. with PBS, 20 *μ*g of H9, 10 *μ*g of IL-2-MSA or 10 *μ*g H9-MSA (generated in the laboratory of Chris Garcia) in a total volume of 100 *μ*l PBS, which was repeated every three days (except for Fig. S1, where it was repeated every two days) until the mice were euthanized or one week after a mouse had no palpable tumor. For the comparison of innate simulants, mice were injected i.t. with PBS or 50 *μ*g of either CDN or CpG ODN DS-L03 (Invivogen) in a total volume of 100 *μ*l PBS.

In some experiments, mice were depleted of NK cells, CD8 cells and/or CD4 cells by i.p. injections of 200 *μ*g of anti-NK1.1 (clone PK136, purified in our laboratory), anti-CD8*β*.2 (clone 53.5.8, Leinco) or anti-CD4 (clone GK1.5, Leinco), respectively, 2 days and 1 day before treatment initiation, and again every six days thereafter until mice were euthanized or no palpable tumor was detected for one week. Whole rat immunoglobulin G (IgG; Jackson ImmunoResearch) was used as a control for depletions. Depletions were confirmed by flow cytometry. In some experiments, mice received 500 *μ*g of anti-IFNAR-1 (clone MAR1-5A3, Leinco) or 200 *μ*g anti-TNF-*α* (clone TN3-19.12, Leinco) i.p. on days -1 and 0 before the initiation of therapy, and again every three days throughout the experiment.

### MCA tumor experiments

Mice were injected subcutaneously on the right flank with 100 *μ*g of 3-methylcholanthrene (Crescent Chemical) in peanut oil, after which they were monitored weekly. When palpable tumors (∼50 mm^3^) were detected, usually 8-10 weeks after MCA injection, the mice were randomized and entered into the study. Tumors were injected i.t. with PBS or 50 *μ*g of CDN in a total volume of 100 *μ*l PBS on days 0, 3, and 6, which was repeated every 6 six days. Mice were also injected intraperitoneally with PBS or 10*μ*g H9-MSA in a total volume of 100 *μ*l PBS starting on day 0, which was repeated every three days. In some experiments, mice were depleted of NK cells, CD8 cells and/or CD4 cells by i.p. injections of 200 *μ*g of anti-NK1.1 (clone PK136), anti-CD8*β*.2 (clone 53.5.8, Leinco) or anti-CD4 (clone GK1.5, Leinco), respectively, 2 days and 1 day before treatment initiation, and again every six days thereafter until mice were sacrificed. In other experiments, mice received additional therapy, in the form of 200 *μ*g anti-PD-1 (clone RMP1-14, Leinco) and/or 200 *μ*g anti-TIGIT (clone MUR10A from Arcus Biosciences or 1G9 from BioXCell) each in a volume of 100 ul PBS, every three days. In most cases the anti-TIGIT was staggered 1 day after anti-PD-1 treatment, but in some cases it was injected on the same day.

### Two-tumor experiments

All mice were depleted of CD4 and CD8 T cells. Tumor cells were washed and resuspended in PBS. 100 *μ*l containing 4 × 10^6^ cells were injected subcutaneously on the right flank of the mice and 100 *μ*l containing 2 × 10^6^ were injected subcutaneously on the left flank of the mice. Tumor growth was measured using digital calipers and tumor volume was estimated using the ellipsoid formula: V = (4/3)*π*abc. Five days after tumor inoculation, when the right flank tumors were ∼50 to 100 mm^3^, the right flank tumors were injected intratumorally with PBS or CDN and the mice were treated i.p. with H9-MSA or PBS as described above for single tumor experiments.

### Toxicity

To assess toxicity of the CDN/H9-MSA treatment, tumors were established, and mice were treated i.t. with PBS or 50 *μ*g CDN and i.p. with PBS or 10 *μ*g or 20 *μ*g H9-MSA. Mouse weights were recorded daily. Pulmonary edema was determined by measurement of lung wet weight/body weight ratios six days after initiating treatment.

### Flow cytometry

Single-cell suspensions of tumors were generated by dicing tumors with a razor blade followed by enzymatic digestion for 30 minutes at 37C in medium containing 1 mg/mL Collagenase D (Roche Diagnostics) and 1*μ*g/mL DNase I (ThermoFisher Scientific). The samples were further dissociated in a gentleMACS Dissociator (Miltenyi) before passage through a 70-*μ*m filter. Cells were stained directly for determining NK cell numbers and Ki67 staining or incubated for 4 hours in medium containing Brefeldin A (Biolegend) and Monensin (Biolegend) for granzyme B staining before surface staining and intracellular staining. LIVE/DEAD stain (ThermoFisher Scientific) was used to exclude dead cells. Before staining with antibodies, FcgRII/III receptors were blocked by resuspending the cells in undiluted culture supernatant of the 2.4G2 hybridoma cell line (prepared in the lab) and incubating for 20 minutes at 4C. Cells were washed in PBS containing 2.5% FCS and stained with antibodies directly conjugated to fluorochromes for 30 minutes at 4C in the same buffer. For intracellular staining of granzyme B and Ki67, cells were fixed and permeabilized using Cytofix/Cytoperm buffer (BD Biosciences) and stained with antibodies directly conjugated to fluorochromes for 1hr at RT in 1X Perm/Wash buffer (BD Biosciences). NK cells were gated as viable, CD45^+^, CD3^-^, CD19^-^, F4/80^-^, Ter119^-^, NK1.1^+^, NKp46^+^ cells. Flow cytometry was performed using an LSRFortessa or an LSRFortessa X-20 (BD Biosciences). Data were analyzed using FlowJo software.

### Antibodies

For flow cytometry, the following antibodies were used (all purchased from BioLegend): anti- CD45 (30-F11, Alexa Fluor 700), anti-CD3*ε* (145-2C11, APC-Cy7, PE-Cy5), anti-CD4 (GK1.5, BV421), anti-CD8a (53-6.7, BV650), anti-CD19 (6D5, PE-Cy5), anti-Ter119 (TER-119, PE- Cy5), anti-F4/80 (BM8, PE-Cy5), anti-NKp46 (29A1.4, PerCP-Cy5.5), anti-NK1.1 (PK136, BV711, FITC), anti-CD69 (H1.2F3, BV605), anti-Sca-1 (D7, BV510), anti-Ki67 (SolA15, eFluor 450), anti-CD107a (1D4B, Alexa Fluor 647), and anti-IFN-*γ* (XMG1.2, PE). Anti-granzyme B (GB11, PE-CF594) was purchased from BD Biosciences.

### Ex vivo cytotoxicity assay

Cytotoxicity by splenocytes was assessed with a standard 5-hour ^51^Cr-release assay. 24 and 72 hours after treating mice with PBS, CDN (i.t.), H9-MSA (i.p.) or CDN/H9-MSA, spleens were harvested and single cell suspensions were treated with ACK lysis buffer. The ^51^Cr-release assay was performed as described (Nicolai et al, 2019) using B16-F10 *B2m*-/- cells as target cells. For NK cell-depleted samples, mice were injected twice (2 days and 1 day before the initiation of treatment) intraperitoneally with 200 *μ*g of anti-NK1.1 (PK136) and spleens were collected and treated as described above. Efficacies of cellular depletions were confirmed by flow cytometry.

### Statistics

Statistics were performed using Prism (GraphPad). For tumor growth and survival data, two-way analysis of variance (ANOVA) and log-rank (Mantel-Cox) tests were used. Two-way ANOVA was used for cytotoxicity. For flow cytometry data, single therapies were compared to either control PBS or CDN/H9-MSA using one-way ANOVA followed by Dunnett’s multiple comparisons. Significance is indicated as follows: *P < 0.05; **P < 0.01; ***P < 0.001; ****P < 0.0001.

## Supplementary Materials

Fig. S1. Synergistic impact of H9 or H9-MSA administration and CDN therapy on tumor rejection.

Fig. S2. Spider plots showing growth of individual tumors from Figure 1.

Fig. S3. Toxicity associated with H9-MSA or combination therapy.

Fig. S4. Verification of *in vivo* cellular depletions.

Fig. S5. Blockade of TNF-*α* and IFNAR-1 partially reverses the tumor growth delay imparted by CDN/H9-MSA administration in mice depleted of T cells and NK cells.

Fig. S6: Spider plots showing growth of individual tumors from Figure 2.

Fig. S7: Spider plots showing growth of individual tumors from Figure 3.

Fig. S8: Spider plots showing growth of individual tumors from Figure 4.

Fig. S9: Additional activation markers of NK cells induced by CDN/H9-MSA treatments of tumors.

Fig. S10: Representative gating strategies for flow cytometry data in Figure 5A.

Fig. S11: Representative gating strategies for flow cytometry data in Figure 5B.

Fig. S12: Spider plots showing growth of individual tumors from Figure 6.

Fig. S13: Combination immunotherapy for treating primary sarcomas induced by the carcinogen methylcholanthrene.

## Acknowledgements

We thank Yeara Jo, Djem Kissiov, Alan Tubbs and Joanna Kritikou for helpful comments on the manuscript, Michel DuPage for helpful discussions, and Erik Seidel and Susanna Dang for technical assistance. We thank Hector Nola and Alma Valeros in the Cancer Research Laboratory at UC Berkeley for expert assistance with flow cytometry and cell sorting.

## Funding

National Institute of Health grant R01AI113041 (D.H.R.)

UC Berkeley Immunotherapeutics and Vaccine Research Initiative supported by Aduro Biotech 045535 and 045538 (D.H.R.)

National Science Foundation predoctoral fellowship, DGE 1752814 (N.K.W.) QB3 Frontiers in Medical Research predoctoral fellowship (N.K.W.)

National Institute of Health predoctoral fellowship, F31CA228381 (C.J.N.) Parker Institute for Cancer Immunotherapy (K.C.G)

Howard Hughes Medical Institute (K.C.G.)

National Institute of Health grant RO1-AI51321 (K.C.G.)

## Author contributions

Conceptualization: NKW, CB, DHR Funding acquisition: DHR, NKW Investigation: NKW, CB, LZ, GS

Methodology: NKW, CB, LP, CJN, KCG, DHR Project administration: DHR

Resources: LP, COH, SMM, KCG Visualization: NKW, DHR

Writing – original draft: NKW, DHR

Writing – review & editing: NKW, CB, LP, CJN, SMM, KCG, DHR

## Data and materials availability

The cyclic dinucleotides used in this study were provided under a material transfer agreement from Aduro Biotech (now Chinook Therapeutics), but molecularly identical CDNs can be purchased. The H9 sequence is published ^35^ and is available to any investigator who would like to generate the protein. All data are available in the main text or the supplementary materials.

## Supplemental Materials

**Figure S1.**
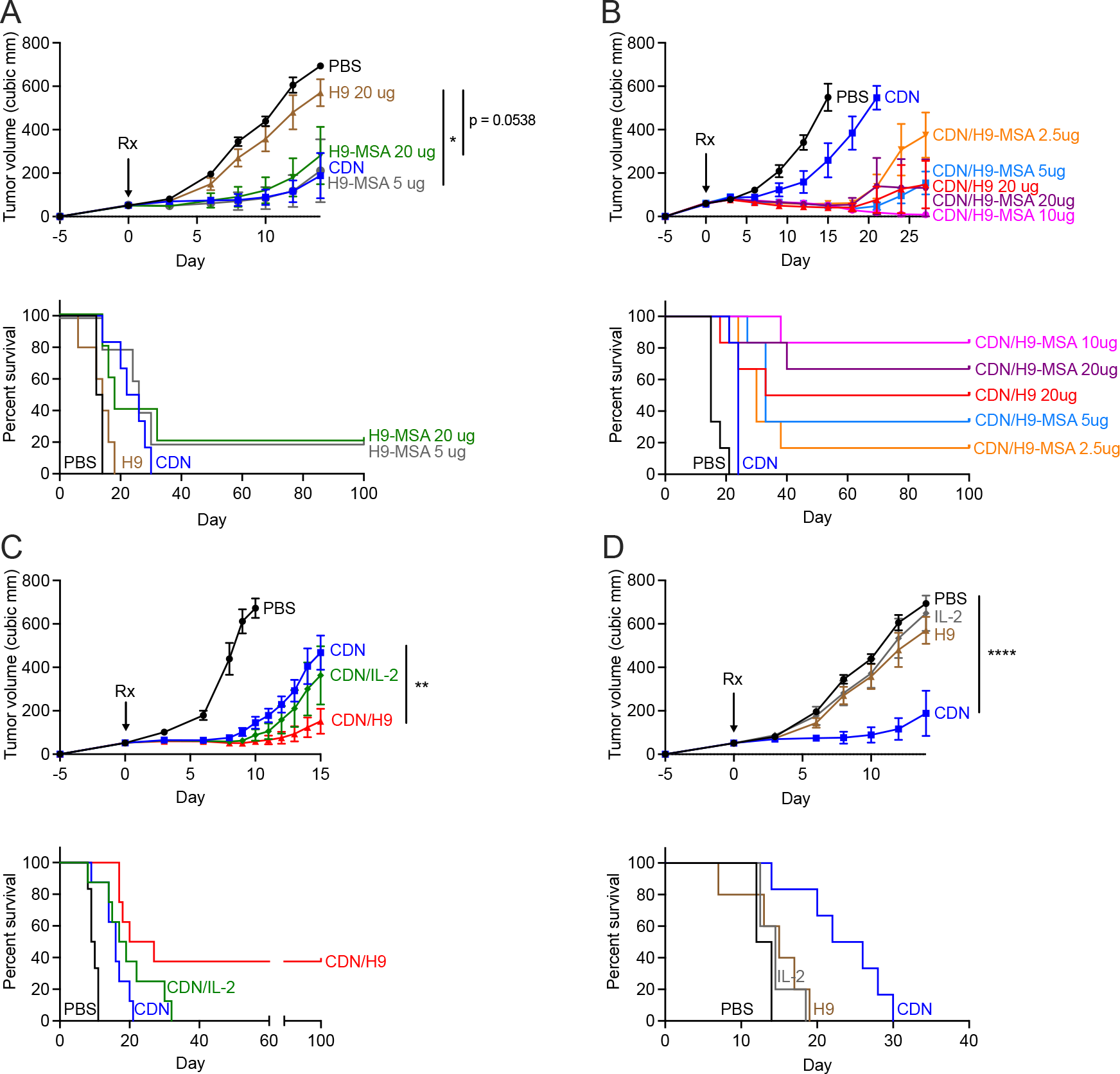
Synergistic impact of H9 or H9-MSA administration and CDN therapy on tumor rejection. B16-F10 *B2m-/-* tumor cells (4 x 10^6^) were implanted s.c.in C57BL/6J mice on day -5 and grown to approximately 50 mm^3^. On d0, tumors were injected once intratumorally with 50 *μ*g CDN or PBS, and/or i.p. with the indicated cytokines or PBS. Cytokine administrations were repeated every two days until mice were euthanized or until 1 week after a mouse completely cleared a tumor. Tumor growth curves and survival data are shown. (A) Half-life extended H9 (H9-MSA) showed significant antitumor effects as a monotherapy, even with a lower administered dose (5 *μ*g). (B) Dose titration of H9-MSA in combination with CDN therapy, showing that doses >5 *μ*g were necessary for optimal combination therapy effects. (C) When combined with CDN therapy, H9 administration, but not IL-2 administration, resulted in improved antitumor effects including long term survival of a significant percentage of the animals. (D) CDN monotherapy, but not monotherapy with IL-2 or H9, delayed tumor growth. The data in panels A-C were from one large experiment. Tumor growth data were analyzed by 2- way ANOVA. Survival data were analyzed using log-rank (Mantel-Cox) tests. The data in all panels are representative of two independent experiments. n=6-7 mice per group.

**Figure S2.**
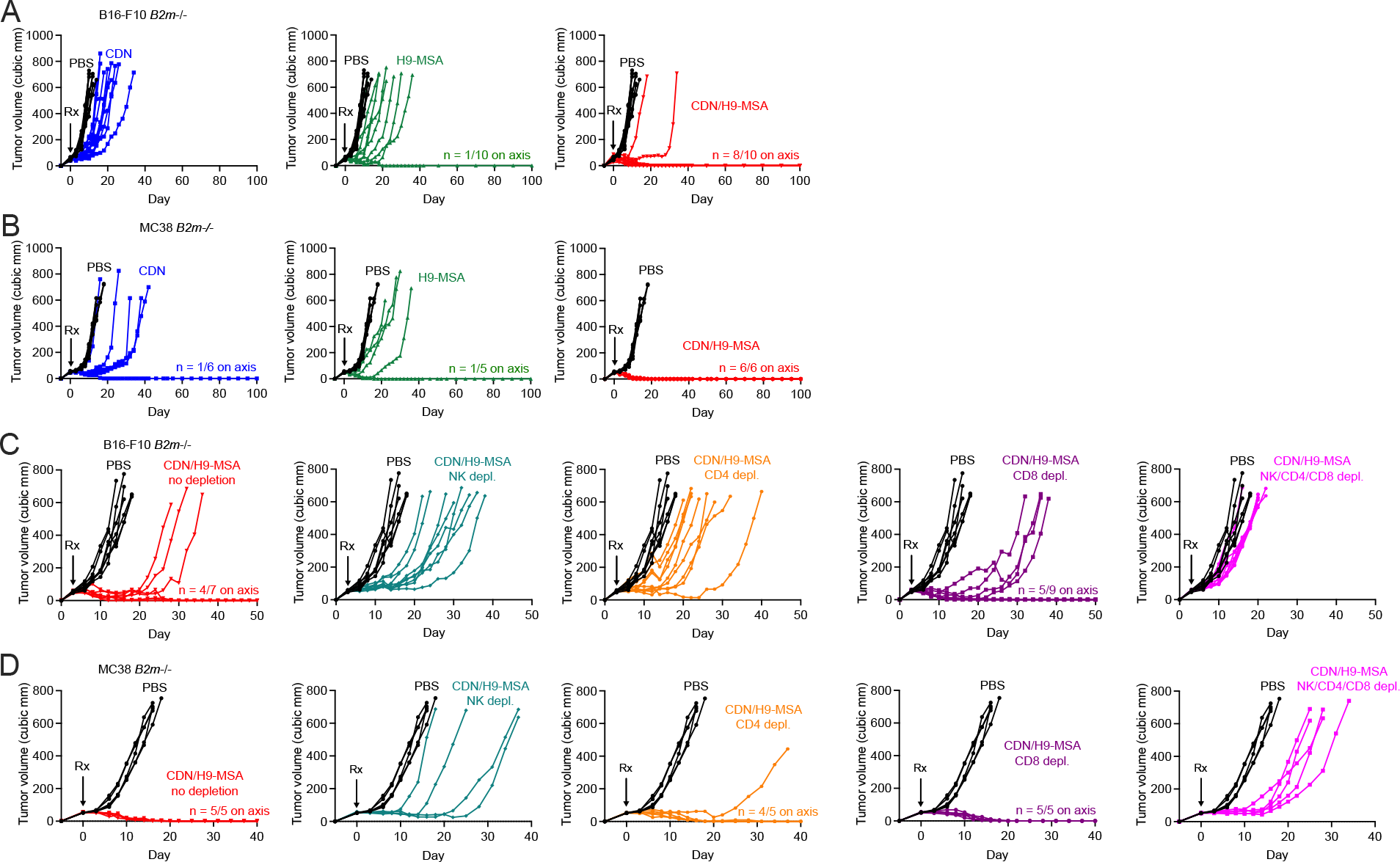
Spider plots showing growth of individual tumors from. Figure 1. **(A, C)** B16-F10 *B2m*-/- tumors **(B, D)** MC38 *B2m*-/- tumors. **(A, B)** impact of combination therapy vs monotherapies. **(C, D)** Impact of cellular depletions on the efficacy of CDN/H9-MSA combination therapy.

**Figure S3.**
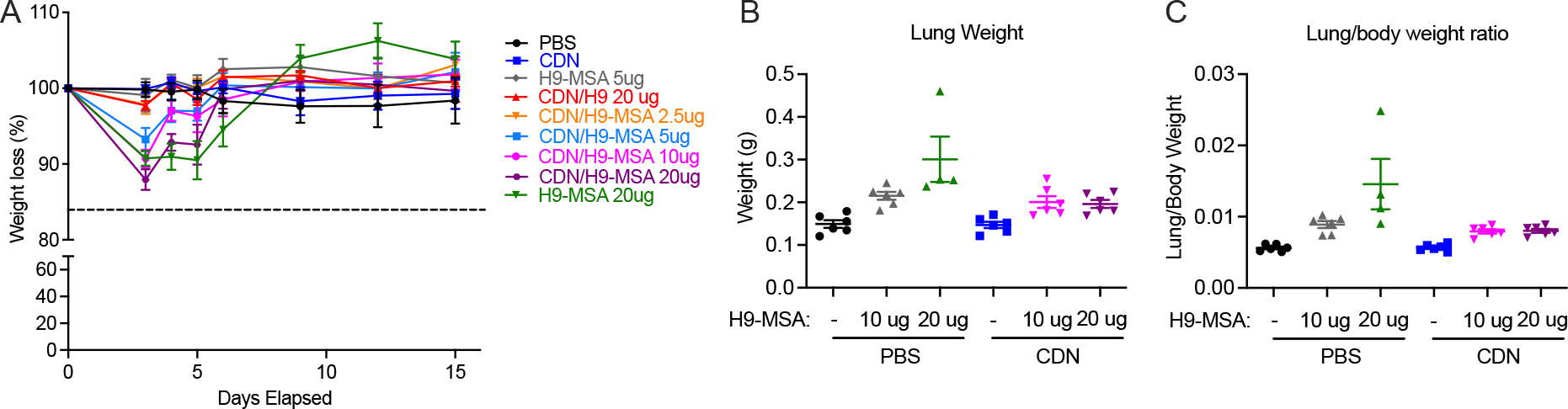
Toxicity associated with H9-MSA or combination therapy. Tumors were generated and mice received therapy as described in Figure S1, legend. Mice were weighed on the indicated days after initiating therapy (d0). In a separated experiment, lungs were harvested on day 6, and wet weights were determined. **(A)** Percentages of d0 body weights over time. **(B)** Lung weight (grams) on day 6. **(C)** The lung/body weight ratio, a measure of pulmonary edema, is shown for each mouse on day 6. Each analysis was confirmed in a repeat experiment.

**Figure S4.**
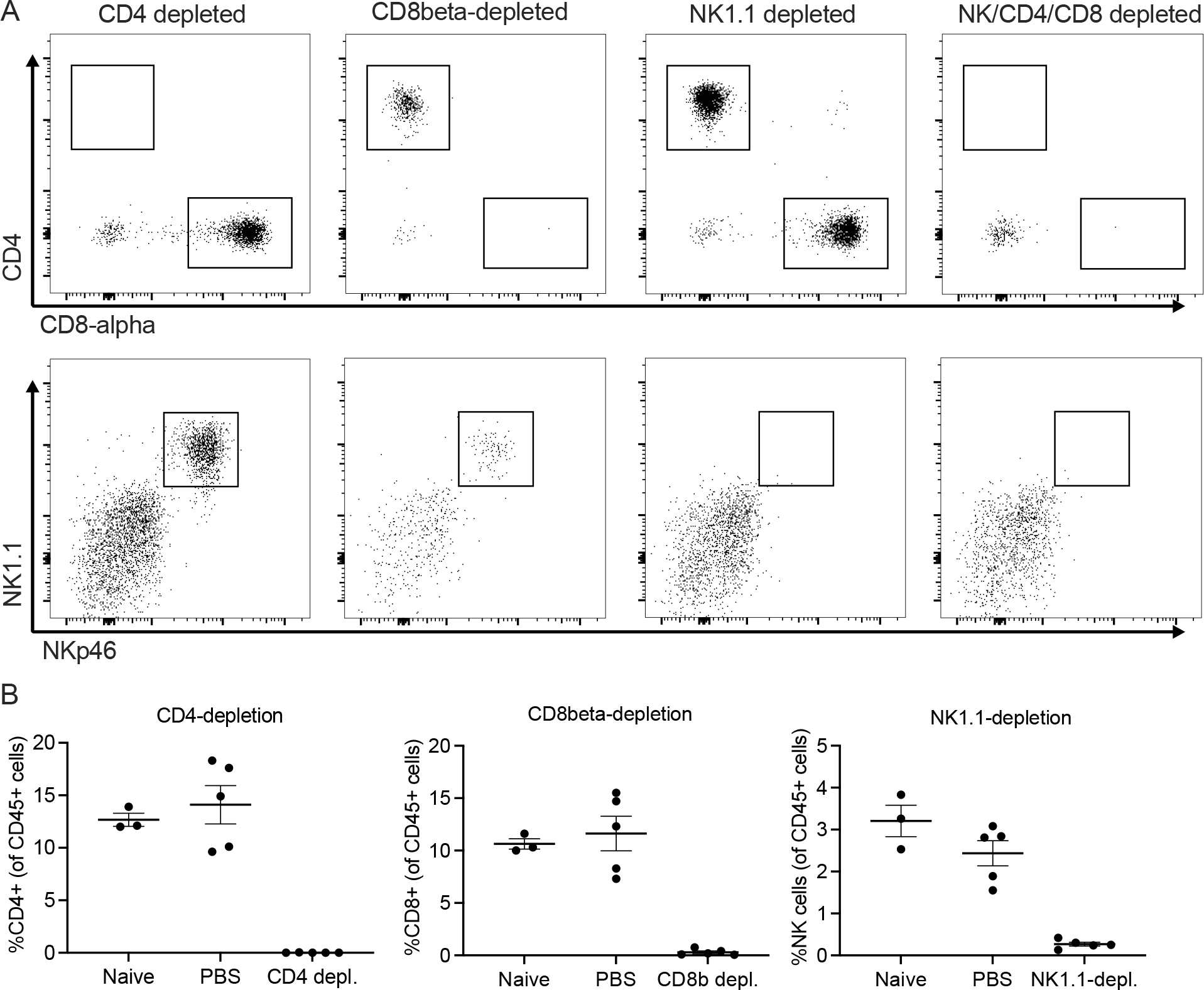
Verification of *in vivo* cellular depletions. **(A)** Successful T cell and NK cell depletion protocols. As described in Methods, mice were treated on days -2 and -1 with antibodies to deplete CD4+ cells, CD8*β*+ cells, NK1.1+ cells or all three. On day 0, blood cells were collected, ACK-treated. Gated viable, CD45+, CD19-, Ter119-, F4/80- cells were examined. **(B)** Representative percentages of the indicated cells after the indicated depletions compared to contemporaneous controls.

**Figure S5.**
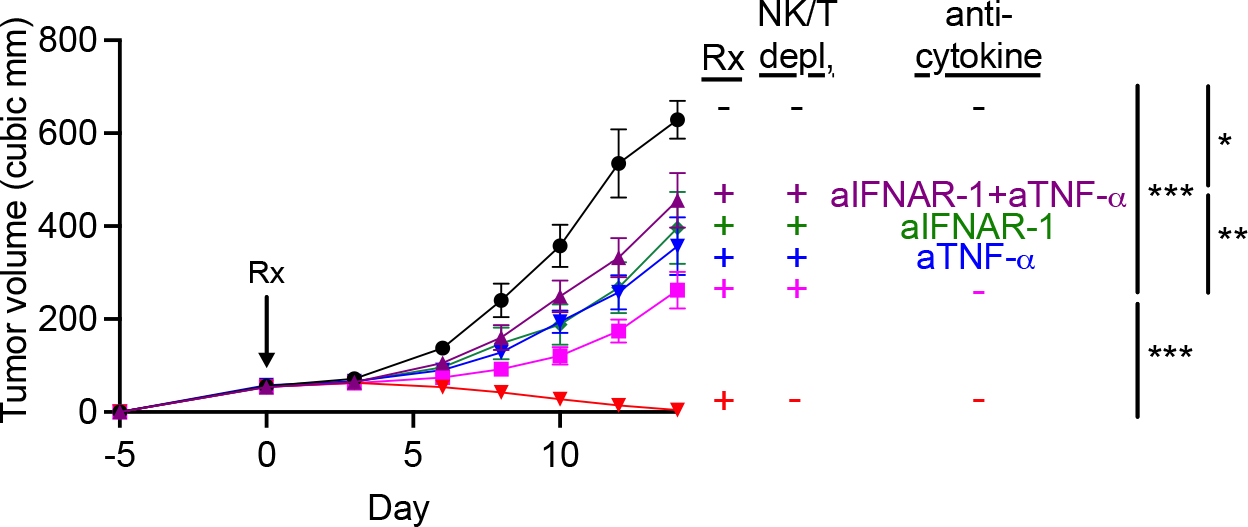
Blockade of TNF-*α* and IFNAR-1 partially reverses the tumor growth delay imparted by CDN/H9-MSA administration in mice depleted of T cells and NK cells. Tumors were generated and mice received therapy as described in Figure S1, legend. Mice were depleted of NK cells, CD4 T cells and CD8 T cells i.p. on days -2 and -1 before the initiation of therapy and repeated every 6 days. Additionally, mice received anti-IFNAR-1 and/or anti-TNF-*α* i.p. on day -1 and day 0 before the initiation of therapy and again every three days. Blockade of TNF-*α* and IFNAR-1 led to more rapid tumor growth in the NK/T cell depleted animals.

**Figure S6:**
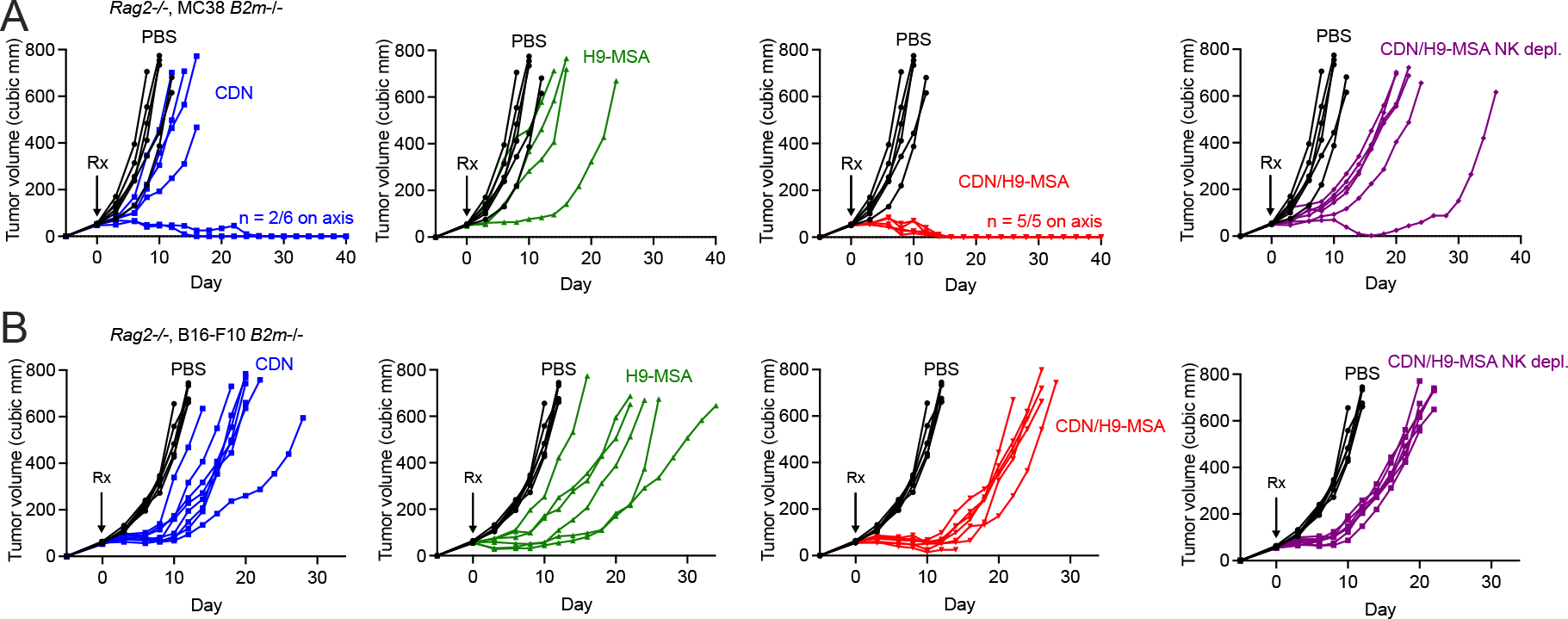
Spider plots showing growth of individual tumors from Figure 2. (A) ***Rag2-/-***mice with MC38 *B2m*-/- tumors. **(B)** *Rag 2-/-* mice with B16-F10 *B2m*-/- tumors.

**Figure S7:**
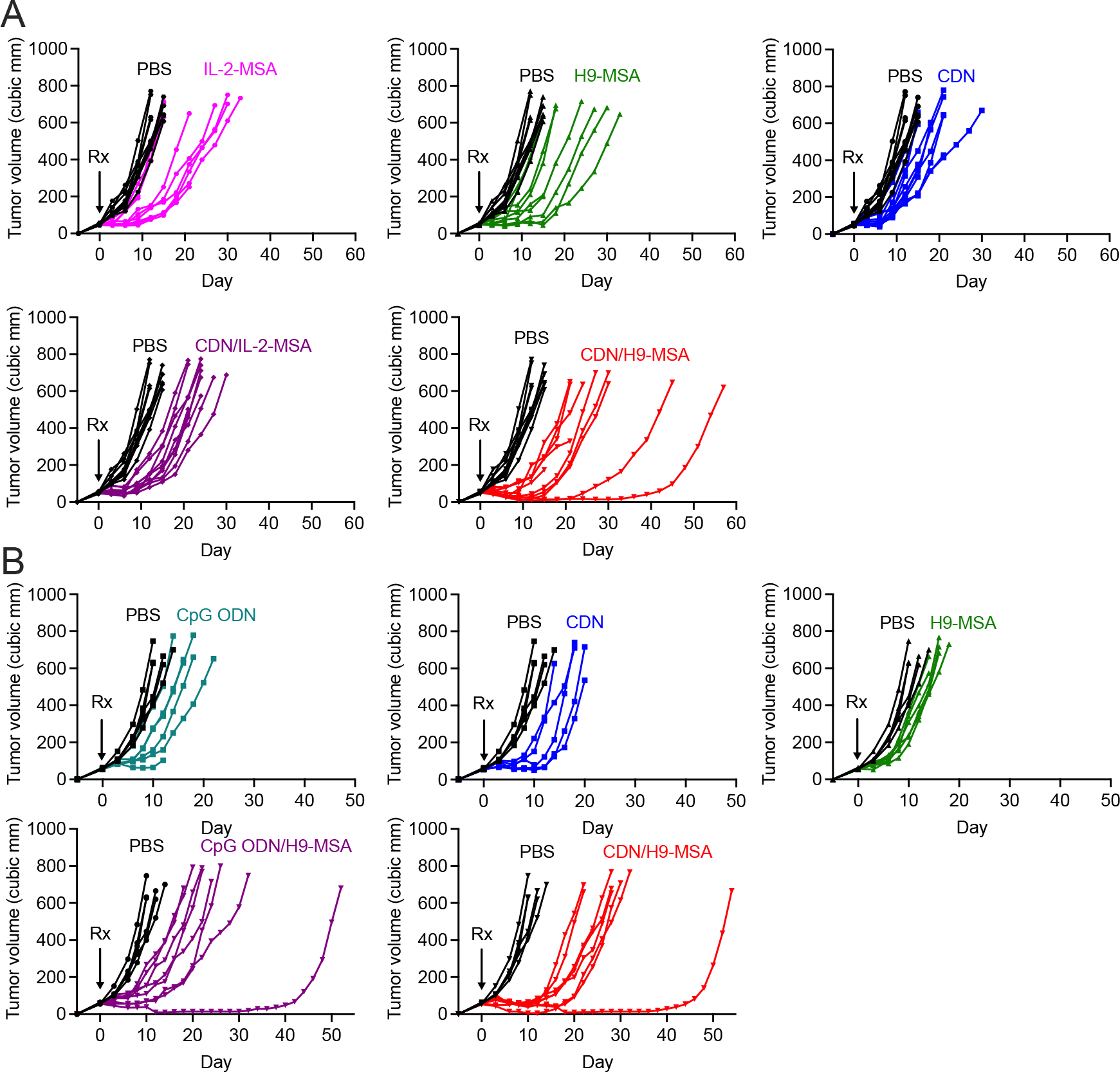
Spider plots showing growth of individual tumors from Figure 3. **(A)** Spider plots for comparison of IL-2-MSA and H9-MSA treatments. **(B)** Spider plots for comparison of CpG ODN and CDN combination treatments.

**Figure S8:**
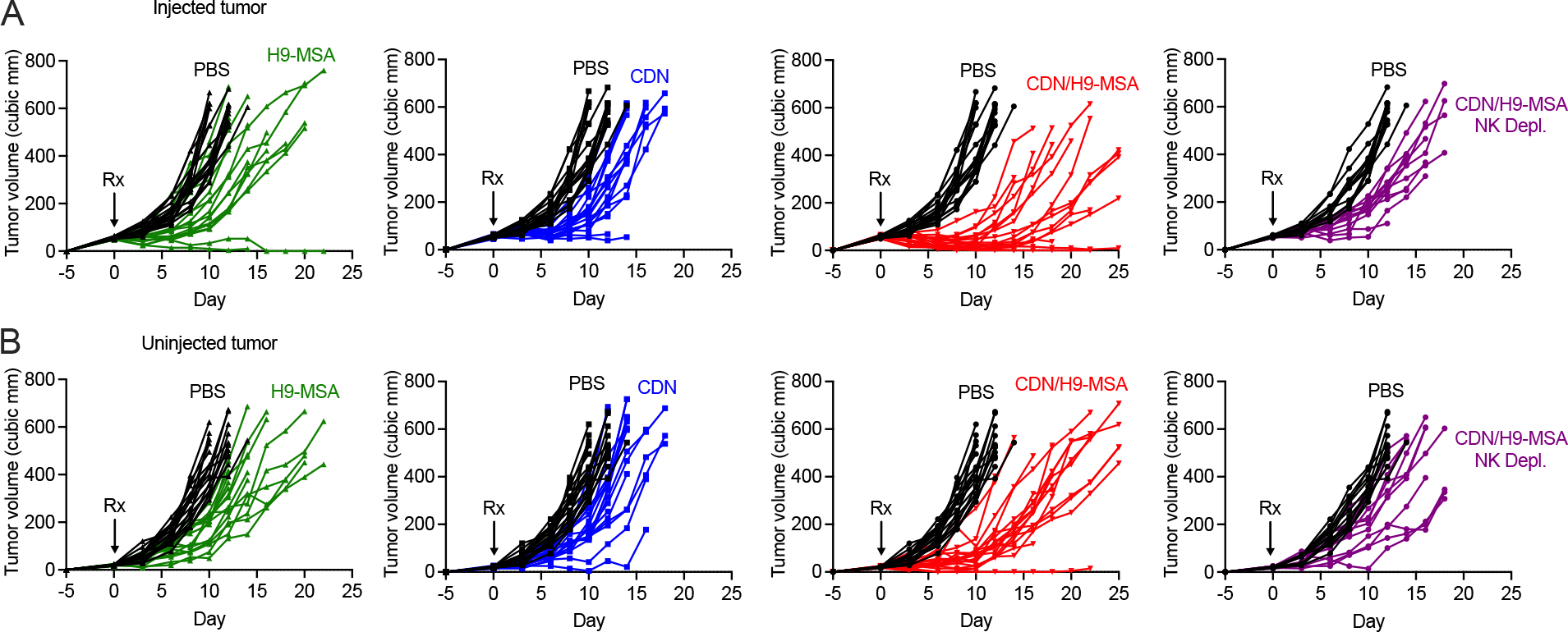
Spider plots showing growth of individual tumors from Figure 4. **(A)** Injected tumors and **(B)** Uninjected tumors from Fig. 4.

**Figure S9:**
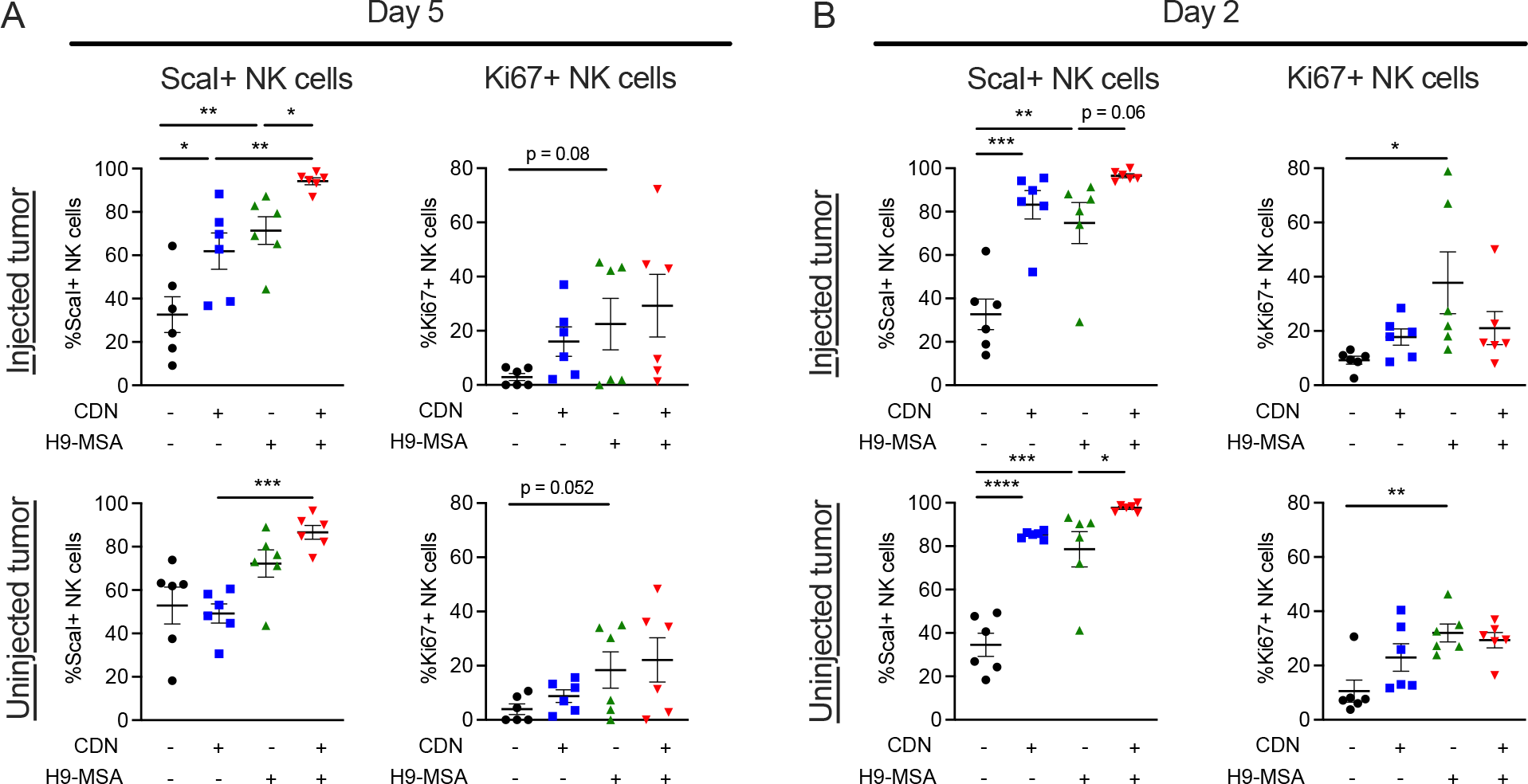
Additional activation markers of NK cells induced by CDN/H9-MSA treatments of tumors. **(A, B)** B16-F10 *B2m*-/- tumors were established on both flanks of C57BL/6J mice, treated and analyzed 5 days (A) and 2 days (B) after initiating therapy as in Figure 5. NK cells were gated as viable, CD45^+^, CD3^-^, CD19^-^, F4/80^-^, Ter119^-^, NK1.1^+^, NKp46^+^ cells. n=3. Data are representative of at least 2 independent experiments.

**Figure S10:**
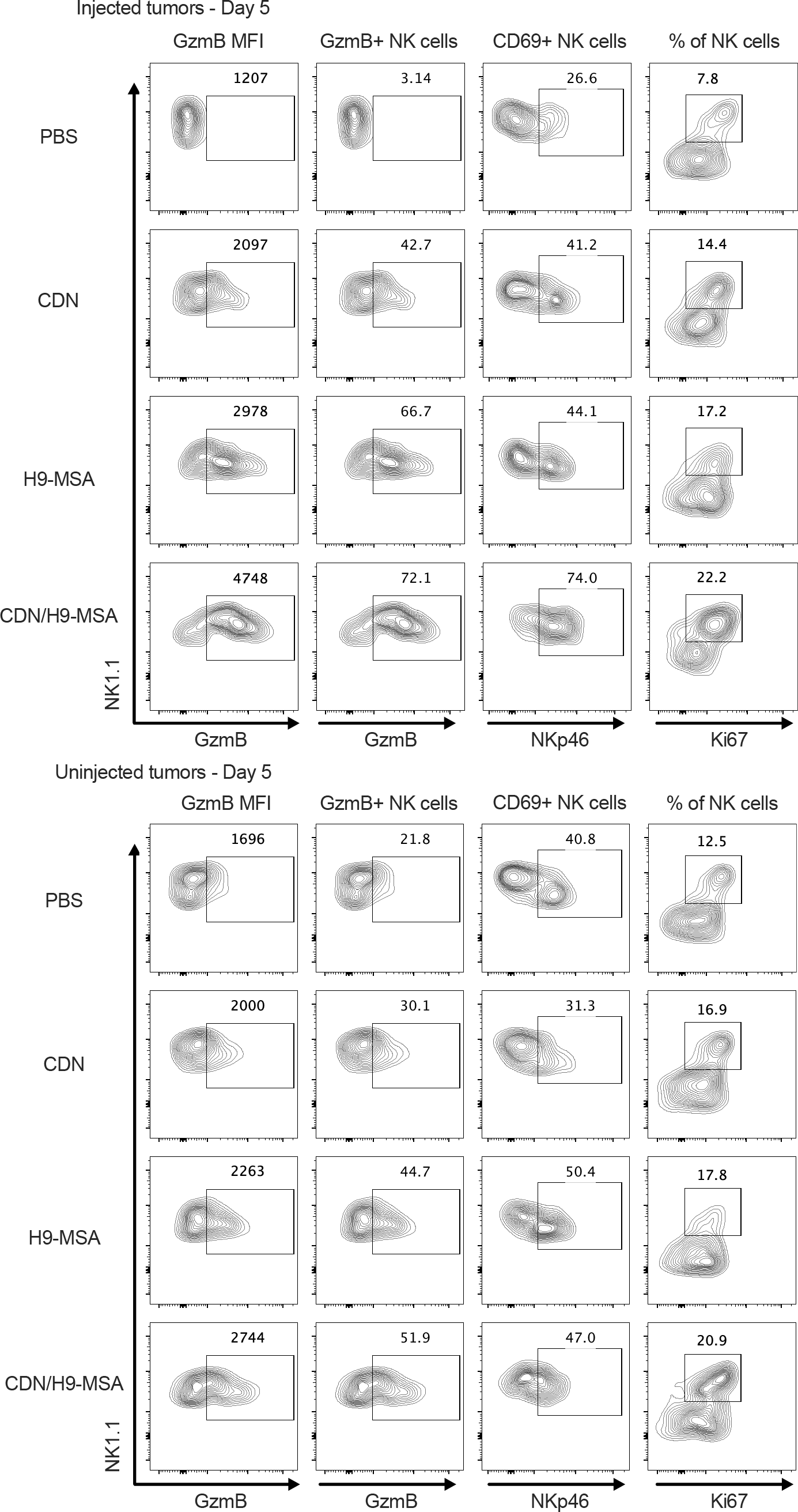
Representative gating strategies for flow cytometry data in Figure 5A. NK cells were gated as viable, CD45^+^, CD3^-^, CD19^-^, F4/80^-^, Ter119^-^, NK1.1^+^, NKp46^+^ cells.

**Figure S11:**
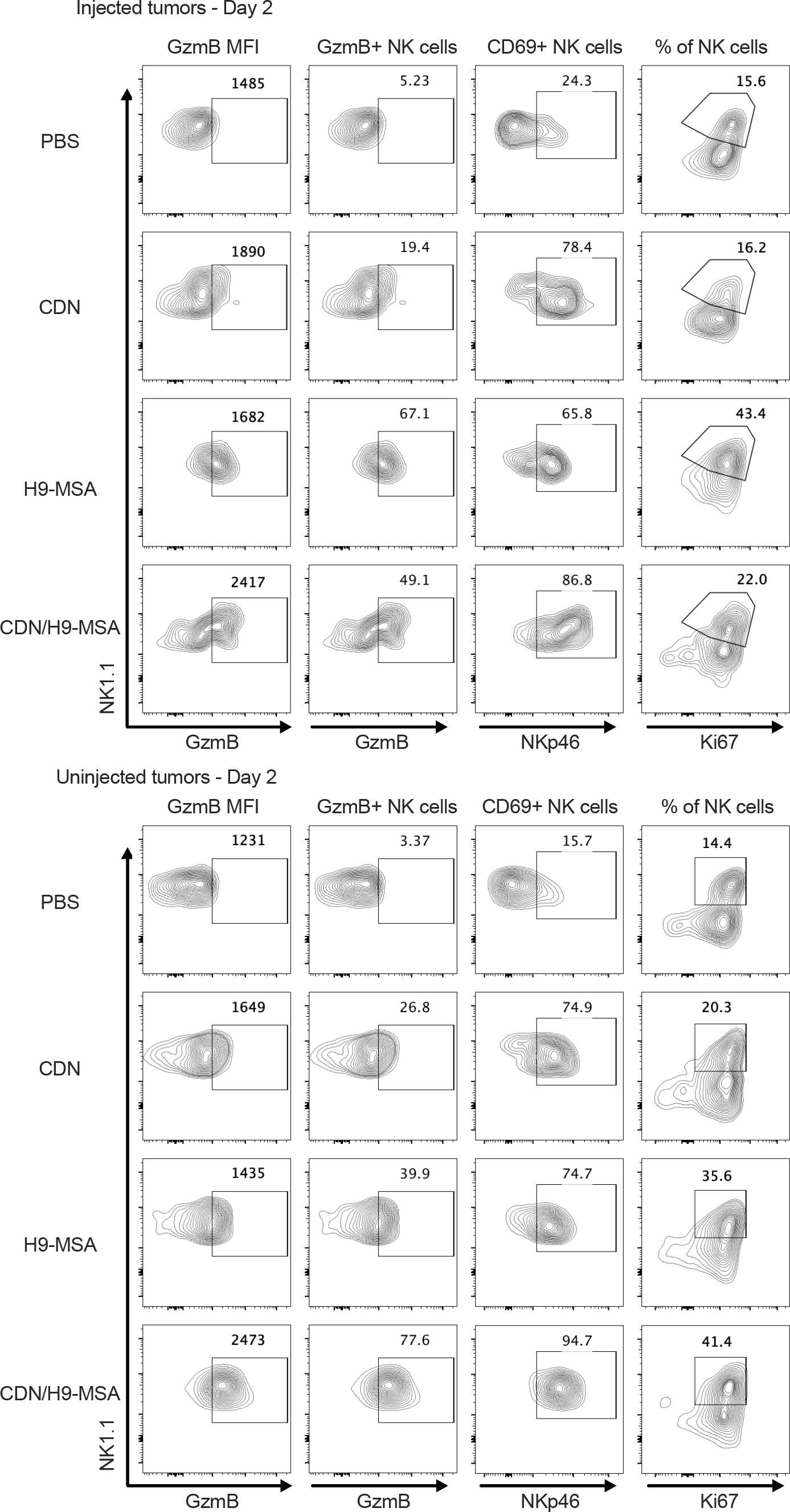
Representative gating strategies for flow cytometry data in Figure 5B. NK cells were gated as viable, CD45^+^, CD3^-^, CD19^-^, F4/80^-^, Ter119^-^, NK1.1^+^, NKp46^+^ cells.

**Figure S12:**
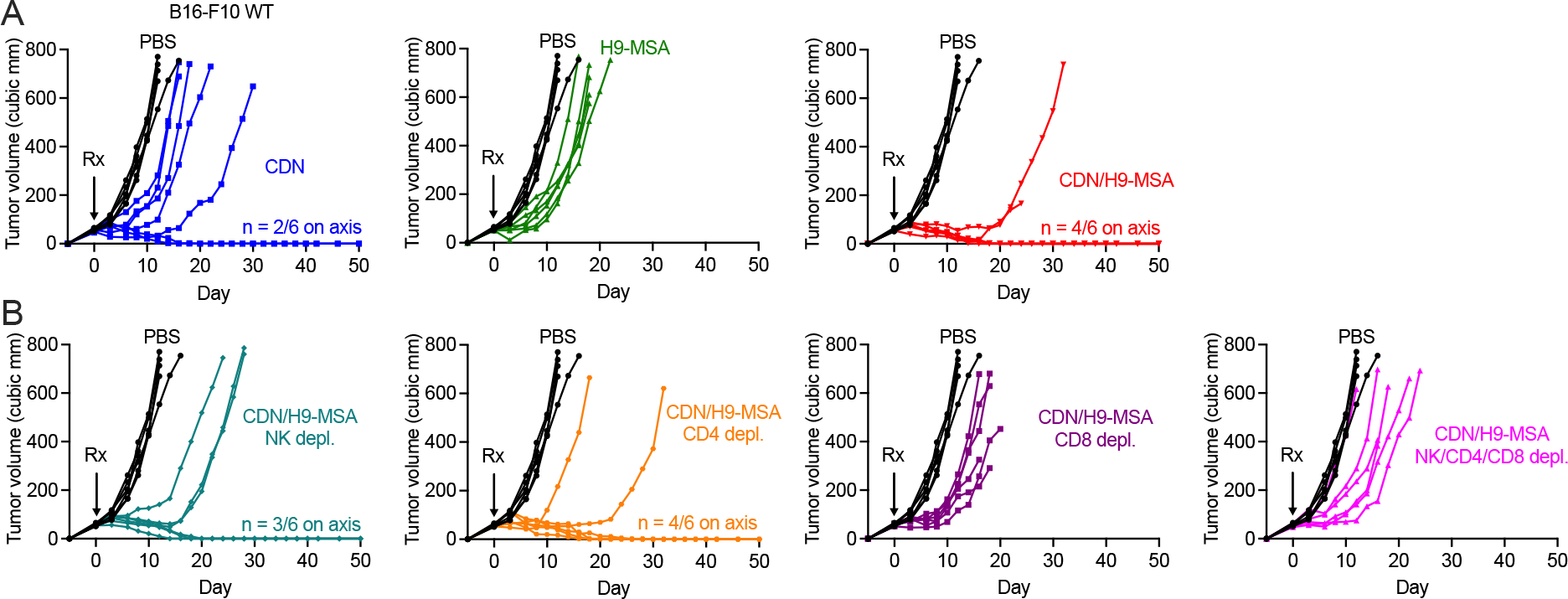
Spider plots showing growth of individual tumors from Figure 6. **(A)** B16-F10 WT tumors and **(B)** B16-F10 WT tumors subjected to cellular depletions.

**Figure S13:**
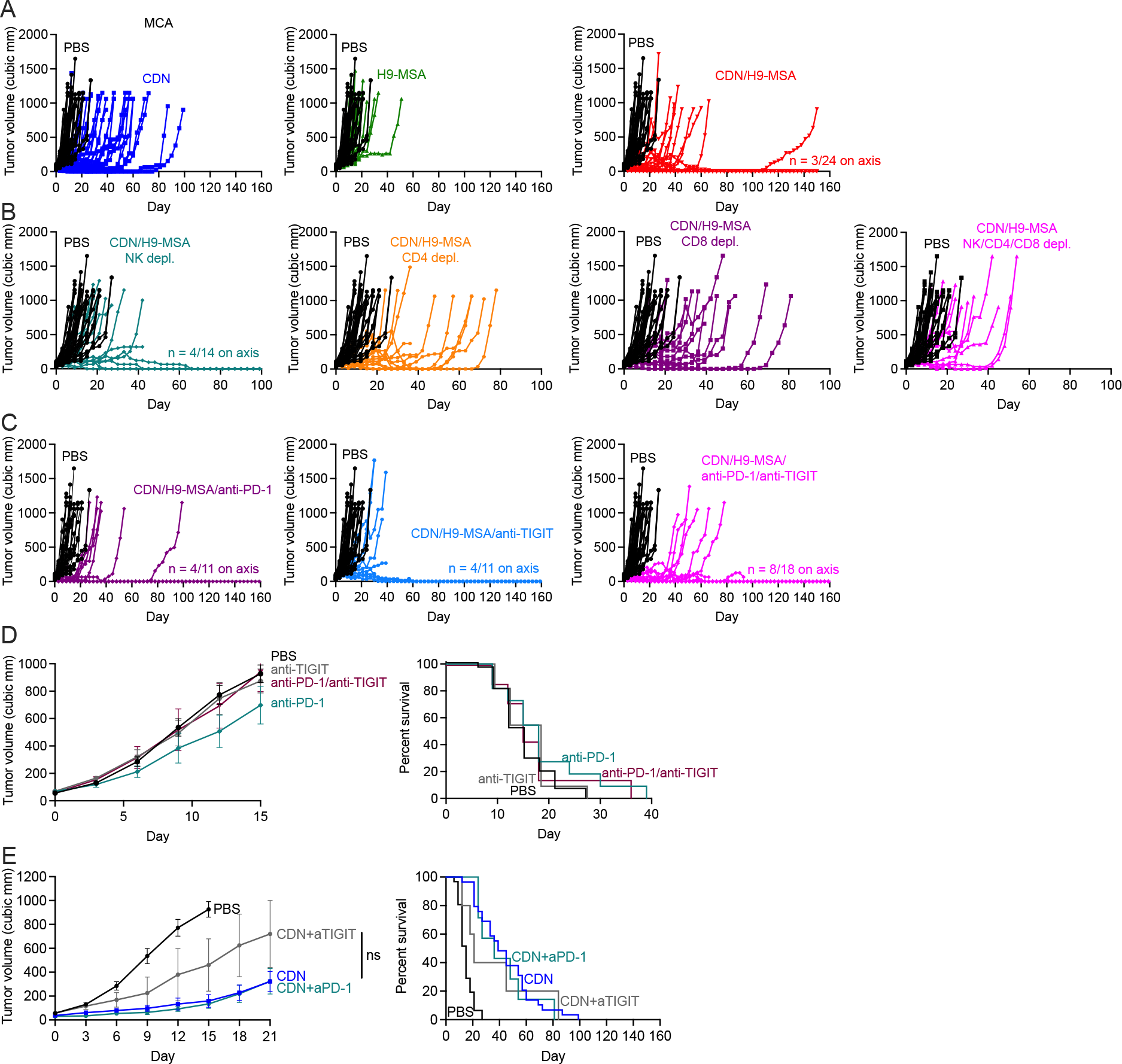
Combination immunotherapy for treating primary sarcomas induced by the carcinogen methylcholanthrene. **(A)** Spider plots showing individual tumor growth for treated MCA induced tumors from Fig. 6D. **(B)** Spider plots for depletions of MCA induced tumors from Fig. 6E. **(C)** Spider plots for CDN/H9-MSA+checkpoint inhibitor combinations from Fig. 6E. **(D, E)** Tumor averages for single agent checkpoint inhibitors **(D)** or checkpoint inhibitors in combination with CDN **(E)**. The anti-TIGIT antibody employed in these panels was clone 1G9 (BioXCell).

